# Dynamics-Informed Priors (DIP) for Neural Mass Modelling

**DOI:** 10.1101/2025.09.26.678721

**Authors:** Alessia Caccamo, Dominic M Dunstan, Mark P Richardson, Alexander D Shaw, Marc Goodfellow

## Abstract

Neural Mass Models (NMMs) are important mathematical tools for inferring hidden neural mechanisms that generate healthy and pathological brain activities. A critical step in the inference process is parameter estimation, which calibrates NMMs based on measured neuroimaging data. While parameter estimation can be conducted via various approaches, one of the most influential methods is Dynamic Causal Modelling (DCM). DCM adopts a Bayesian inference approach that relies on, and is sensitive to, the specification of prior parameter distributions reflecting *a priori* hypotheses about the causes of data. However, most parameters of NMMs encode neuronal properties that are not directly measurable. For this reason, in the absence of sufficient empirical data and well-founded prior beliefs, inference becomes increasingly susceptible to bias. Therefore, it was imperative to establish a comprehensive strategy for mapping model parameters to data. This study proposes a computational extension of DCM, named DCM with dynamics-informed priors (DIP-DCM), which adopts a genetic algorithm (GA) to map parameter values to model dynamics. Optimal sub-regions of the parameter space were subsequently selected and translated into groups of parameter priors for DCM. DIP-DCM was compared to the ‘standard’ DCM inference and to the standalone GA, using two independent neuroimaging datasets. Results indicated that DIP-DCM models were the best predictors of data and captured key mechanistic signatures of psychiatric disease and pharmacological interventions. Overall, DIP-DCM addressed degeneracy, handled local minima, and explored diverse parameter regimes following trajectories informed directly by model dynamics and data. This study suggests that DIP-DCM is an advantageous route to parameter estimation when information is limited, enabling a data-driven derivation of parameter priors – or hypotheses – in exploratory studies, across different biological contexts and datasets.

## 1 Introduction

Large-scale cortical activity is widely recognised to emerge from the functional integration of distinct and spatio-temporally distributed neural events (Schomer & da Silva, 2017). While the causal relationships between these events remain inherently hidden and inaccessible to direct empirical observation, their collective effects manifest as signals measured via neuroimaging methods, such as electroencephalography (EEG), magnetoencephalography (MEG) and functional magnetic resonance imaging. Tools from dynamical systems theory, statistical inference, and computational analysis have become invaluable for mapping hidden neuronal processes and interactions to observable signals. Specifically, Neural Mass Models (NMMs), with their long history dating back to 1938 (de Ńo, 1938; Freeman, 1975; Lopes da Silva et al., 1974; Mountcastle, 1957), are often utilised to simulate cortical dynamics and to study emergent neural population phenomena. NMMs are abstract representations of local brain circuits and treat neural populations as single entities, or point processes. Therefore, neural population activity is described by average membrane potential and firing rate variables, and the population parameters reflect *lumped* neural properties, including synaptic strength, response timescales and rates of activity propagation. These models enable causal inference, the process that identifies the causes – encoded by the respective biophysical parameters – that underlie the observed empirical data and its modulation across different states of healthy functioning or neurological disease (David & Friston, 2003; Glomb et al., 2022).

However, the problem of reconstructing causes from observed effects, i.e., the inverse problem, does not generally offer a unique solution: multiple parameter regimes, under the same model, can generate the same data (Dunstan et al., 2023; Hartoyo et al., 2019; Hashemi et al., 2025). This behaviour is often referred to as parameter “non-identifiability” (Chis et al., 2016; White et al., 2016) or “degeneracy”, and it is an important ubiquitous property of real biological systems (Edelman & Gally, 2001; Marder & Goaillard, 2006; Prinz et al., 2004). Degeneracy refers to the presence of different mechanistic pathways for the same biological function and it is an inevitable consequence of natural selection. Notably, this ‘many-to-one’ relationship contributes to the resilience and stability of biological functions, yet, it becomes problematic in the context of mechanistic modelling. Here, degeneracy can be influenced by several aspects including the model structure, the data used in the model fitting process, the number of free parameters and the parameter estimation strategy. The latter is of notable importance as parameter inference can be conducted using different techniques, and the chosen methodology is likely to affect the mapping between parameter values and observed data.

Among the existing methods, Dynamic Causal Modelling (DCM) (Friston et al., 2003) provides a well-established and principled framework for parameter inference in neuroimaging. Within DCM, model parameters are estimated using a computationally efficient variational Bayesian routine, known as Variational Laplace (VL) (Friston et al., 2003; Parr et al. T. & Friston, 2022; Zeidman et al., 2023). Data are modelled using a generative framework that defines prior expectations, or beliefs, about their underlying causes, along with a likelihood function that describes the probabilistic relationship between data and causes (Friston et al., 2003; Lee & Vanpaemel, 2018). Through a process of belief updating, parameter priors – the probability distributions assigned to parameters *a priori* – are updated into posterior distributions, used to derive the most probable explanations for the observed data. Therefore, parameter priors are a crucial requisite for solving the DCM inverse problem.

However, as the inverse problem can display a ‘many-to-one’ relationship between hidden causes and their manifestation, the DCM VL is predisposed to settle on one solution among many options, as predetermined by the chosen parameter priors (Morita et al., 2010). Identifying the most suitable priors, however, can be challenging. For instance, in the context of NMMs, parameters are abstract representations of real neural population properties and do not directly correspond to measurable quantities (Coombes, 2023; Deschle et al., 2021; Dunstan et al., 2023; Ferrat et al., 2018; Goodfellow et al., 2022). Similarly, for novel models or datasets it can be difficult to quantify and establish reasonable and objective beliefs about the model parameters. While these issues can be mitigated by conducting systematic explorations of the prior parameter space, these can become inefficient when parameter spaces are vast, and ineffective when the explored priors do not reflect the ‘true’ generative process.

Previous endeavours adopted alternative optimisation techniques for DCM, including a gradient-free Markov Chain Monte Carlo (MCMC) sampler and a gaussian process optimisation (GPO), which used GPO-identified solutions to initialise a local gradient descent (Lomakina et al., 2015; Sengupta et al., 2015). Specifically, the GPO method improved parameter estimation accuracy compared to traditional variational Bayes, and enhanced computational efficiency compared to MCMC. However, this method was limited to providing information on the mode of the posterior distribution rather than approximating a posterior density. In addition, its application was restricted to hemodynamic equations and required sufficiently smooth and convex objective functions, i.e., the mathematical expressions that guide parameter estimation and quantify model performance. Thus, the GPO approach was considered less suitable for models of electrophysiological data (Lomakina et al., 2015), since they are more likely to display non-convexity and hence to be characterised by multiple local minima. Local minima are suboptimal solutions, or solutions that optimise the objective function strictly relative to nearby points. They are commonly encountered in the context of non-smooth objective functions, which are highly sensitive to small parameter perturbations. This sensitivity introduces irregularities, or ‘fractures’, in the parameter landscape, complicating the identification of global optima. Therefore, there is a need for novel strategies that guarantee robust parameter estimation, not only by ruling out poorly informed priors, but also by mitigating challenges posed by such complex relationships between objective functions and parameter values.

To address these challenges, the present study proposes a novel method which derives parameter priors directly from data by leveraging heuristic decision processes to shape and constrain subsequent DCM inversions. This method, termed DCM with dynamics-informed priors (DIP-DCM), adopted genetic algorithms (GAs) to identify multiple parameter configurations to be used as priors within the generative model space. These were named dynamics-informed priors (DIP), as they were shaped by, and learned from, the predicted observations generated by the model under different parameters. Notably, GAs are well-established global optimisation techniques employing the Darwinian principles of natural selection and evolution (Holland, 1992), thus offering a compelling conceptual link between evolutionary selection and selection of generative models. Moreover, GAs were previously found to be useful for parameter estimation in NMMs (Dunstan et al., 2023).

This paper defines the methodological foundations of DIP-DCM and offers two distinct applications through the use of published M/EEG datasets (Biondi et al., 2022; Shaw et al., 2020). Ultimately, this approach can be utilised as an alternative strategy for choosing parameter priors for dynamic causal models, thereby broadening the applicability of DCM to a wider range of exploratory studies. This implementation is compatible with the DCM SPM12 environment and its use is suggested for DCM studies lacking strong prior hypotheses or theoretical assumptions about model parameters, or studies that would benefit from exploring broader parameter regimes at reasonable computational costs.

## 2 Methods

### 2.1 Datasets

#### The neuroimaging datasets were from two independent studies

*Study 1*, Biondi et al., 2022. EEG data were recorded from 14 healthy participants using 64 electrodes. Recordings were performed in resting eyes-closed conditions, at baseline and following the administration of two anti-seizure medications (ASMs), lamotrigine (LTG) and levetiracetam (LEV), or placebo (PL). Recordings were conducted across three days of testing, with drug washout periods of ∼1-2 weeks, resulting in six sub-datasets, or experimental conditions, encompassing all participants (preand post-LTG, LEV and PL). The EEG time series were segmented into 2-second epochs, with the number of epochs ranging from 6 to 72, depending on the participant. The time series were re-referenced to the common average and power spectral densities (PSDs) were computed for each epoch via Fast Fourier Transform for frequency bins from 2 to 45 Hz with 0.5 Hz resolution, as indicated in Biondi et al., 2022. PSDs were normalised to unit area, and averaged across epochs, channels and subjects. The log-transformed grand average PSDs were used as the observed data in the modelling process.

*Study 2*, Shaw et al., 2020. Two hundred and seventy-five channel cryogenic MEG was recorded from patients with schizophrenia and healthy controls during a visual task known to induce gamma oscillations. Data were obtained using the LCMV beamformer on a 4 mm grid. A virtual electrode was derived from the voxel in the visual cortex showing the biggest amplitude change in the gamma range between stimulus ‘on’ and ‘off’ conditions. Data included a total of 5498 and 6268 4-second epochs for patients and controls, respectively. PSDs were computed using Welch’s method (MATLAB pwelch) for frequency bins from 2 to 85 Hz with a Hamming window of 512 points and 50% overlap. PSDs were normalised to unit area, averaged across all epochs and log-transformed.

### 2.2 Neural Mass Model

This study aimed to introduce a novel strategy for selecting parameter priors within the DCM framework as implemented in SPM12. To achieve this, a default SPM12 model was chosen. This was the “LFP” NMM `(spm_fx_lfp` function), that was considered appropriate for both datasets (**Methods 2.1**). Indeed, as reported in the published literature, the “LFP” model can generate a large repertoire of dynamics including high-frequency oscillations – compared to the traditional Jansen and Rit NMM from which it was derived (Jansen & Rit, 1995) – and it is suitable for modelling drug effects (Moran et al., 2007, 2013). Briefly, the model comprises three neuronal populations, or masses, assigned to specific cortical layers and interconnected via synaptic interactions. As usual for NMMs, the activity of each mass is simplified to average post-synaptic potentials and firing rates. The post-synaptic potential produced in response to pre-synaptic firing is modelled by a second-order differential equation, which conveys a linear representation of synapses (‘pulse-to-wave’ or rate-to-potential conversion). The average firing rates are calculated from the post-synaptic potential as the output of a nonlinear ‘wave-to-pulse’ sigmoid function. The model is described in depth in Moran et al., 2007 and in **Supplementary 1**, including details on parameters, their values and biological meanings.

### 2.3 Parameter estimation

For each dataset and related experimental conditions, parameter estimation was conducted via three distinct approaches. (1) DCM, using default SPM12 settings. (2) A GA, which was adapted from Dunstan et al., 2023 and revised for SPM compatibility and for facilitating its integration with DCM models. (3) A novel hybrid combination of GA and DCM, named DCM with dynamics-informed priors (DIP-DCM). DIP-DCM will be the principal focus of this foundational paper.

As is customary in DCM, generative models were specified as a likelihood function and parameter priors. The likelihood function, which remained unchanged across all three approaches, reflected the observed PSD of the model, which was determined by the transfer function of the system under simplifying assumptions (**Supplementary 1**). The priors were the “standard” model priors, as defined in Moran et al., 2007. Parameters were log-scaled to ensure positivity (Friston et al., 2019) and specified as mean and variance of a Gaussian distribution. Parameter estimation was conducted via variational inversion, i.e., variational Laplace (VL), by maximising the negative (variational) free energy (Zeidman et al., 2023). This was defined as the difference between model complexity and accuracy (Friston et al., 2006, **Supplementary 2**).

The GA framework utilised the NSGA-II variant from the MATLAB gamultiobj function (Deb, 2001). Differently from DCM, it searched within closed parameter intervals, spanning biologically realistic values (**Supplementary 1**), rather than adopting parameterised probability densities. Details of the GA are provided in **Supplementary 2**. Briefly, parameter optimisation was conducted by adopting selection, crossover and mutation operators that allow the parameter values to evolve across generations. Parameters were optimised by minimising two objective functions, *J*_1_ and *J*_2_, tailored to the observation of two peaks in the PSDs. The objectives measured the root mean square error (RMSE) between the power spectrum of the model and of the data for different PSD frequencies:

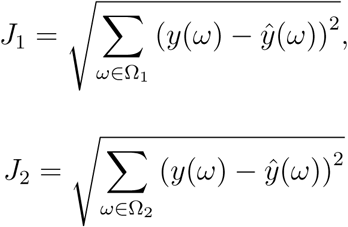

where *ŷ* is the PSD of the model, *y* is the PSD of the data, *ω* is the frequency, and Ω_1_, Ω_2_ are two different frequency ranges, informed by and adapted from empirical observations. Given that the two experiments yielded qualitatively different spectra, the chosen frequencies were Ω_1_ = [6.5, 12.5] Hz, Ω_2_ = [11.5, 30] Hz for study 1 (Biondi et al., 2022) and Ω_1_ = [10.7, 65.8] Hz, Ω_2_ = [35.6, 85] Hz for study 2 (Shaw et al., 2020). Notably, the frequency contributions are allowed to be naturally balanced by the evolutionary processes of the algorithm, bypassing the need for frequency-specific weights, which may not always be generalisable or easy to define *a priori*. However, it is also possible for users to define custom objective functions tailored to specific domain requirements.

The GA yielded sets of parameters, analogous to maximum likelihood estimates, which were used to generate a Pareto front of solutions, i.e., a set of points in the 2D objective space where no solution can be improved without worsening the other. From the Pareto front one solution was selected, i.e., the solution with smallest Euclidean distance between the two objectives in the objective space. The GA was repeated 1000 times under randomised initial conditions sampled via Latin Hypercube (LH) within the specified bounds (Dunstan et al., 2023, 2025).

The DIP-DCM approach combined GA and DCM into a ‘two-steps’ strategy. First, the GA explored the parameter space globally, offering a mapping between parameter values and model dynamics. Second, DCM was used for local ‘exploitation’ of the neighbourhood of points close to the parameter estimate identified by GA. Specifically, among the 1000 GA parameter sets, the top *m* sets were selected based on their model dynamics, as scored by RMSE. Each parameter in the set was used as the mean of a Gaussian distribution that assumed unit variance (i.e., all diagonal elements of the covariance matrix were equal to one), as all estimates were considered equally plausible. These parameter distributions were the dynamics-informed priors defining *m* distinct generative models. Each model entered the VL routine to reach a stable prediction, recovering *m* variational posteriors and the associated free energy values. The *m* posterior densities were formalised into a single distribution as indicated in **Appendix A**, which is a process typically known as Bayesian model averaging (Penny et al., 2010). Notably, by averaging across multiple realisations (at least 400), DIP-DCM reduces the influence of noise and idiosyncratic fluctuations, contributing to more precise predictions as demonstrated in **Supplementary 3**. Nevertheless, since the number of selected priors and inversions is a tunable parameter within this framework, users can tailor the number of priors or inversions based on the specific goals and constraints of their study. Study 1 utilised *m* = 500 and study 2 utilised *m* = 400. A summary of DIP-DCM is provided in **Figure 1** and further details are in **Appendix A**.

**Figure 1:**
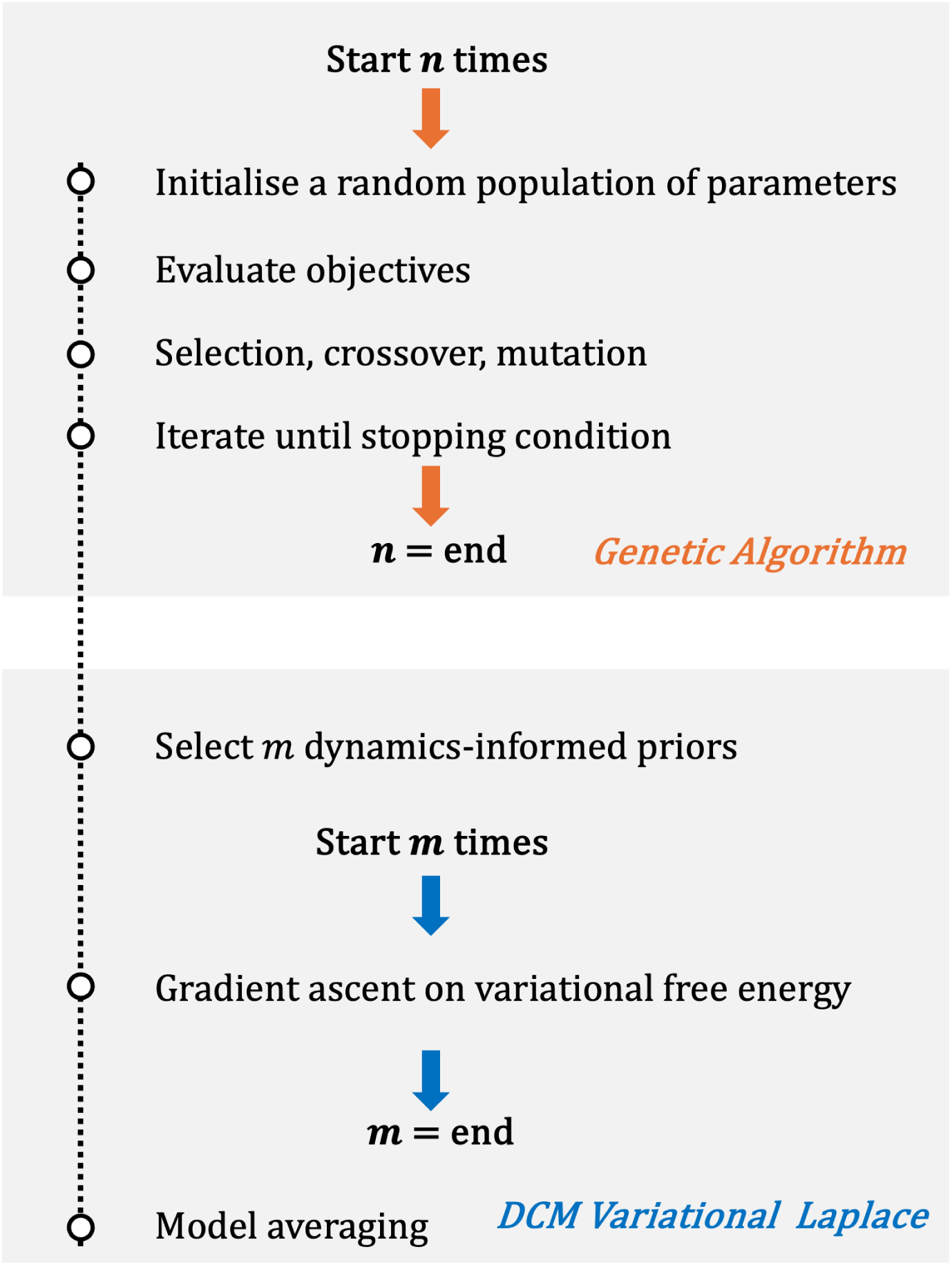
DIP-DCM flow chart. Dynamic Causal Modelling with Dynamics-Informed Priors (DIP-DCM) involves two steps, described in **Methods 2.3**. First, a genetic algorithm is utilised to select appropriate dynamics. Second, the information learned is applied to construct parameter priors, which enter the DCM variational inference routine. *Abbreviations*: GA, genetic algorithm; DCM, dynamic causal modelling.

### 2.4 Inference on the effects of treatments and pathology

Inference was conducted at the group level, and all calculations are formally defined in **Appendix B**. Group-level effects, with groups being pre-treatment and post-treatment in study 1 or control and schizophrenia in study 2, were estimated as the mean absolute difference between 10^5^ draws from the corresponding posterior parameter distributions.

Plausibility of nonzero differences were inferred from the 95% Bayesian Credible Intervals (BCIs) of the difference distributions. Intervals that excluded zero were interpreted as analogous to statistical significance at the *α* = 0.05 level.

These effects were streamlined by interpreting their practical importance on the basis of Cohen’s *d* standardised effect sizes (Sullivan & Feinn, 2012), with standardisation enabling direct mapping between studies, contexts and measurement scales. As convention, *d* = 0.2 indicated a non-trivially small effect, *d* = 0.5 a medium effect, and *d* = 0.8 a strong effect. Within this context, *d* was indicative of the effect magnitude of pharmacological intervention or pathology.

All work was performed using MATLAB (Mathworks Ltd, USA, R2023b) and the SPM12 toolbox.

## 3 Results

To illustrate the challenges associated with parameter estimation, free energy landscapes were derived as indicated in **Appendix C**. **Figure 2** provides a specific example of the effects of parameter priors on the search for the most plausible DCM. This example explores the range of plausible prior means for parameter *G*1, which represents the magnitude of connectivity between pyramidal and stellate cell populations of the model. Notably, this is a *lumped* parameter that would be difficult to measure or calculate directly in real biological systems. Therefore, choosing an accurate prior is also difficult. The “LFP” NMM (**Methods 2.2**; Moran et al., 2007) was fitted to the PSD of EEG data recorded in baseline conditions (pre-placebo dataset from study 1, **Methods 2.1**). The mean of the parameter distribution was varied in log space in the range [−1, 0.85], which included the “standard” prior, log (*G*1) = 0 (Moran et al., 2007, **Appendix C**) and the remaining model parameters were set to their “standard” prior. Variational free energy (**Methods 2.3**) was calculated for each parameter setting via the standard variational inversion, and the information was utilised to construct a free energy profile, as shown in **Figure 2**. Small perturbations in the parameter value were accompanied by changes in the model dynamics (**Figure 2**, between *a*_1_ and *a*_2_), that created discontinuities in the free energy landscape and isolated basins of convergence (sets of points that would converge to the local or global optimum within the range explored). Notably, when the inversion was initiated from the standard prior, the posterior estimate of *G*1 did not correspond to the global maximum free energy, which remained unexplored and required an alternative prior distribution. This illustrates that, depending on the choice of parameter prior, variational inference methods relying on gradient descent techniques can be susceptible to converging on suboptimal solutions, despite the existence of superior solutions elsewhere in the parameter space. These findings highlight a fundamental epistemological implication about the way in which knowledge and beliefs about parameters are acquired and resulting inferences justified. By taking a data-driven approach and exploiting evolutionary processes, DIP-DCM could offer an effective pathway for identifying optimal parameter priors, as addressed in the following sections.

**Figure 2:**
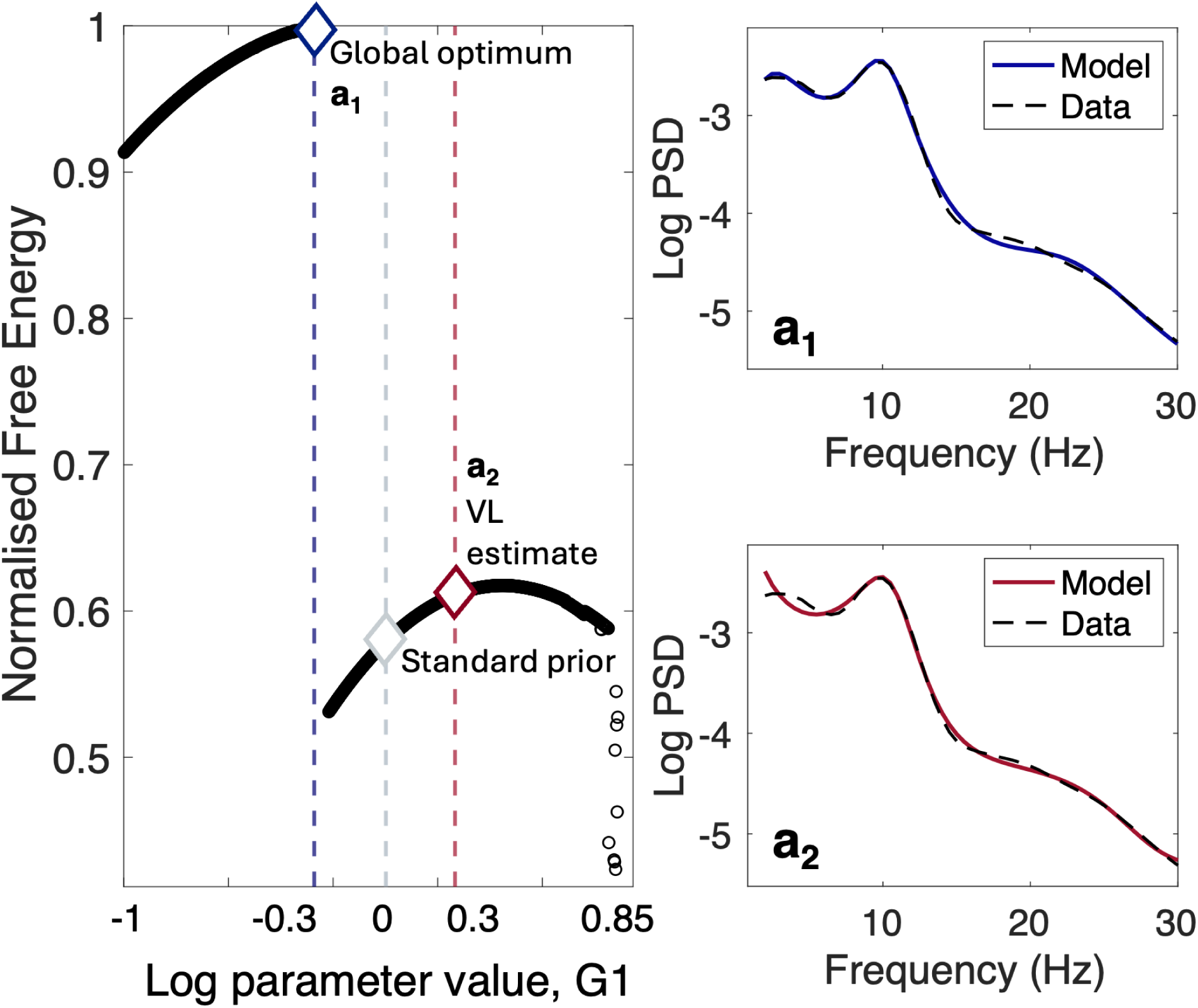
Free energy profile of a connectivity parameter. The mean of the Gaussian distribution of parameter *G*1 (Moran et al., 2007 model) was varied in log space across the biologically plausible range [−1, 0.85]. All remaining parameters were set to the “standard” priors (**Supplementary 1, Table S1**). The standard DCM variational inversion estimated the parameter posteriors and the associated variational free energy for each prior setting (**Supplementary 1**). Variational free energy was normalised between 0 and 1. *a*_1_*, a*_2_: spectra of model corresponding to the colour-coded parameter value, plotted against the data (Biondi et al., 2022, pre-placebo). The global optimum is in blue and the posterior estimate updated from the “standard” prior is in red.

### 3.1 Study 1

#### 3.1.1 Dynamics-informed priors outperform standard priors, yielding more plausible generative models

The dataset (**Methods 2.1**) was collected in healthy participants at baseline and following the administration of LTG, LEV or placebo. Compared to placebo, exposure to LTG and LEV increased the spectral power in the beta band (13-30 Hz; LTG vs placebo, p-value = 0.014 and LEV vs placebo, p-value = 0.011, by Mann-Whitney U-test; **Figure 3A,B**), consistent with prior research on drug-näıve patients with epilepsy and other conditions (Cho et al., 2012; Dini et al., 2023). For each experimental condition (**Methods 2.1**), PSDs were modelled via GA, DCM, and DIP-DCM (**Methods 2.3**) to identify the mechanisms underlying the increase in beta oscillations following ASM administration. For these comparisons, the acronym DCM is used to refer to a DCM adopting “standard” priors.

**Figure 3:**
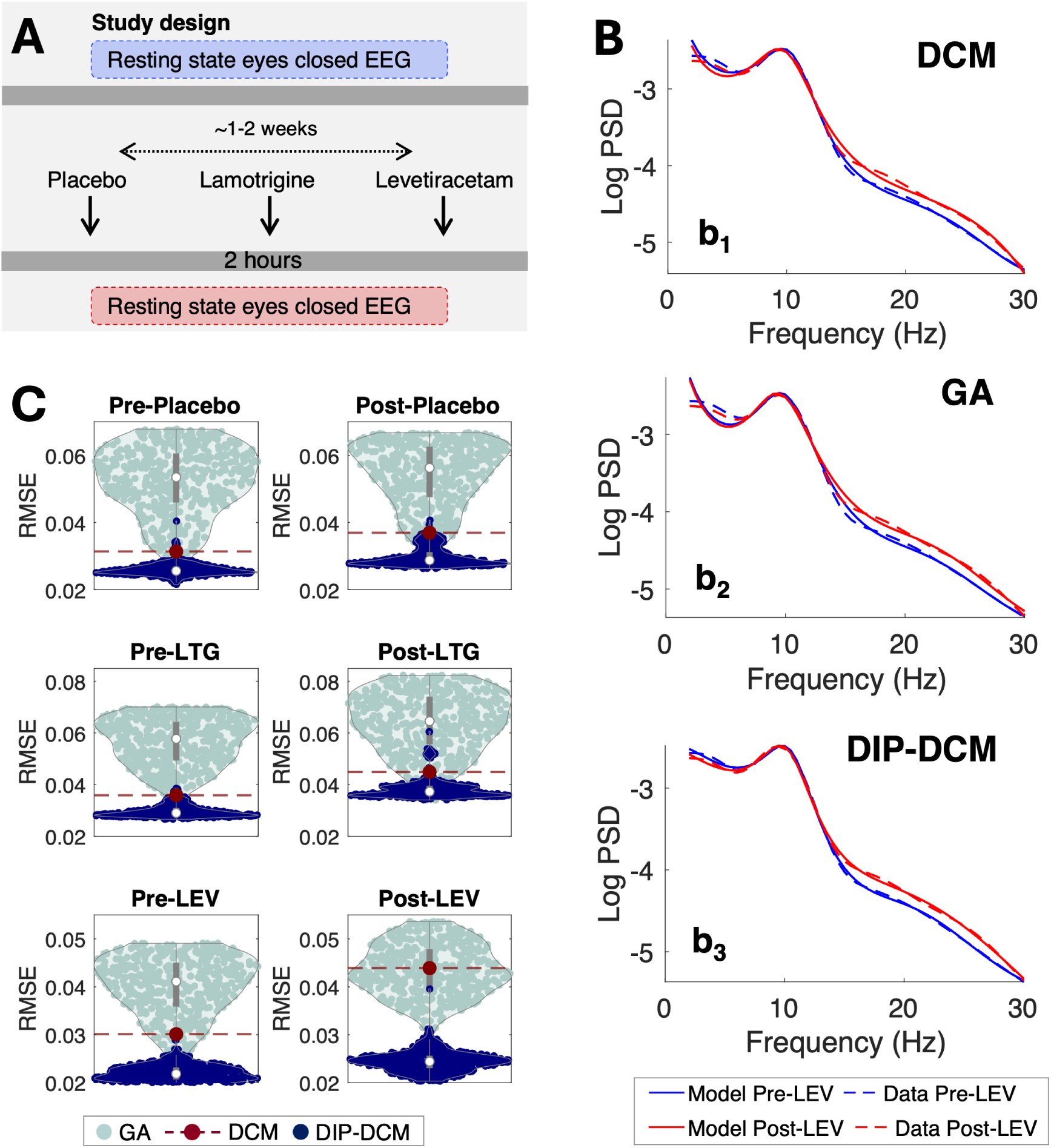
DIP-DCM consistently outperforms GA and DCM. Spectra were calculated from the Biondi et al., 2022 dataset (**Methods 2.1**). **A**: Study design. **B**: Average data and model generated via DCM with “standard” priors (*b*_1_), GA (*b*_2_) and DIP-DCM (*b*_3_). **C**: RMSE between data and models generated by DCM (red, n=1), GA (turquoise, n=500), and DIP-DCM (navy blue, n=500). RMSE was calculated between 6.5 and 30 Hz, in line with the GA objective functions (**Methods 2.3**). *Abbreviations*: DIP-DCM, dynamic causal modelling with dynamics-informed priors; DCM, dynamic causal modelling; GA, genetic algorithm; RMSE, root mean square error; PL, placebo; LTG, lamotrigine; LEV, levetiracetam.

The performance of GA, DCM and DIP-DCM was first evaluated by calculating the root mean square error (RMSE) between model and respective data (**Figure 3B**). The GA produced parameterisations associated with a broad range of dynamics as evidenced by the RMSE values (**Figure 3C**). Some of these dynamics were similar to those of DCM (**Figure 3B,C**, *b*_1_, *b*_2_). However, 87.3% of these were inferior to those of DCM. Crucially, combining GA with DCM via DIP-DCM improved all model calibrations (**Figure 3B**, *b*_3_), with 99.7% improvement from GA (decrease in paired RMSE data from turquoise to blue, **Figure 3C**) and from DCM alone (burgundy, **Figure 3C**). This result indicates that the GA may identify suboptimal parameters, which however lie in the basin of convergence of the global optimum of the free energy landscape. Therefore, GA-derived parameters are best used as informative priors for VL within DIP-DCM, amplifying the advantages of the two methodologies.

#### 3.1.2 DIP-DCM generates reliable parameter inferences

DCM, GA and DIP-DCM generated different parameter distributions. Exemplar posterior parameter distributions generated by the three parameter estimation approaches (**Methods 2.3**) are shown in **Figure 4A**. For specific parameters, such as Te, both GA and DIP-DCM retrieved multi-modal distributions (**Figure 4A**) that display more than one peak compared to DCM (as it assumes normality). Multi-modality can introduce interpretational complexity, yet it also provides a richer understanding of the alternative parameter regimes, or generative mechanisms, supported by the same data. Thus, DIP-DCM can give information on the most dominant parameter values, but it can also highlight alternative parameter trajectories.

**Figure 4:**
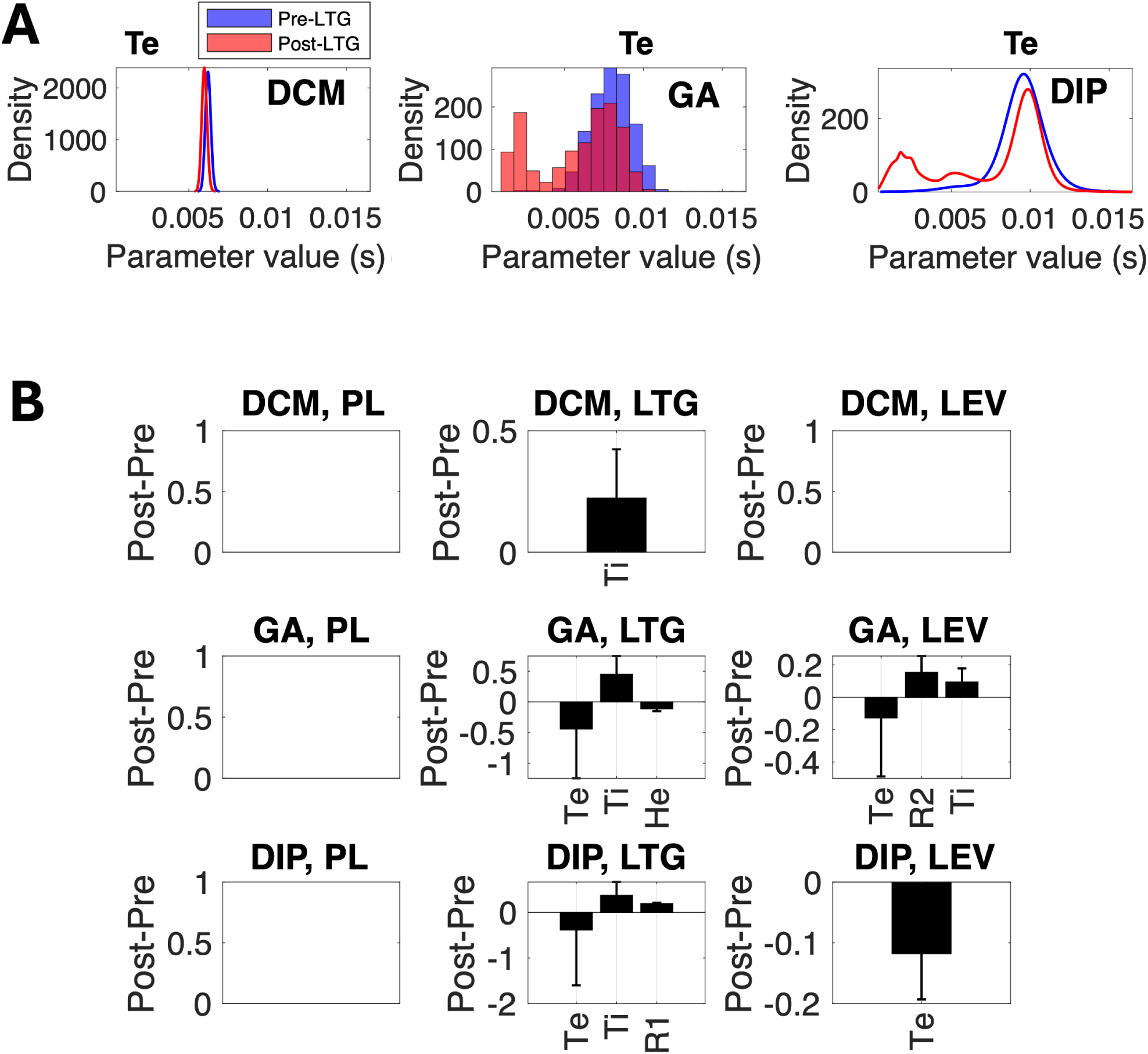
DIP-DCM generates inferences on ASM mechanisms with superior specificity and sensitivity. **A**: Representative examples of posterior parameter distributions for the excitatory time constant parameter Te. All parameter densities are normalised to 1 and displayed within the same range. **B**: Mechanistic effects obtained as per **Methods 2.4**. *Abbreviations*: DCM, dynamic causal modelling; GA, genetic algorithm; DIP-DCM, dynamic causal modelling with dynamics-informed priors; PL, placebo; LTG, lamotrigine; LEV, levetiracetam.

The posterior parameter distributions were used to make mechanistic inferences and derive the effects of placebo, LTG and LEV, as indicated in **Methods 2.4**. Effects were calculated as the absolute difference between mean parameter values at baseline and following pharmacological intervention. Subsequently, effects were selected based on 95% BCI and by considering a sufficient statistical power (Cohen’s d ≥ 0.2). These effects are reported in **Figure 4B**.

While all methods consistently indicated that placebo does not produce detectable effects (**Figure 4B**), discrepancies emerged with respect to the ASMs. DCM predicted a single LTG effect, namely, an increase in the inhibitory synaptic time constant parameter (Ti). Conversely, GA and DIP-DCM both identified an increase in Ti but also a decrease in the parameter Te, representing the excitatory synaptic time constant, and an increase in parameter R1. Moreover, DCM did not predict any LEV effects, similarly to the placebo condition, despite the observed increase in beta power (**Figure 3B**). On the contrary, GA revealed an increase in Ti and a decrease in Te, while DIP-DCM identified a single LEV-specific effect, i.e. a selective decrease in Te. These effects are discussed in **Discussion 4.1-2**.

#### 3.1.3 DIP-DCM provides the best trade-off between reliable inference and computational efficiency

A potential drawback of global search heuristics such as GA is that they can be computationally expensive compared to the DCM variational inference. Notably, DIP-DCM optimised the trade-off between accuracy and computational cost, outperforming the standalone GA (**Figure 5A**). The performance of DIP-DCM was further assessed as indicated and shown extensively in **Supplementary 3**. In the first step of DIP-DCM parameters evolve over a number of generations set to be 500. Therefore, this number was reduced to determine the minimum required to generate models with suitable dynamics. Results suggested that a minimum of 150 generations was necessary to maintain a stable model fitness as scored by RMSE, while maintaining computation times moderate (**Figure 5A**). Computation times were ≈ 1 min for 1 full DIP-DCM inversion, or optimisation, while a standard DCM could be inverted in ≈ 30 seconds (data not shown), on a personal computer. However, when iterating over a large number of priors, models were run on a server with 48 cores, or on a supercomputer (ISCA, University of Exeter) that enables efficient parallelisation. Therefore, computation times can differ depending on the machine, on the parallelisation approach, and on the number of priors selected by the user.

**Figure 5:**
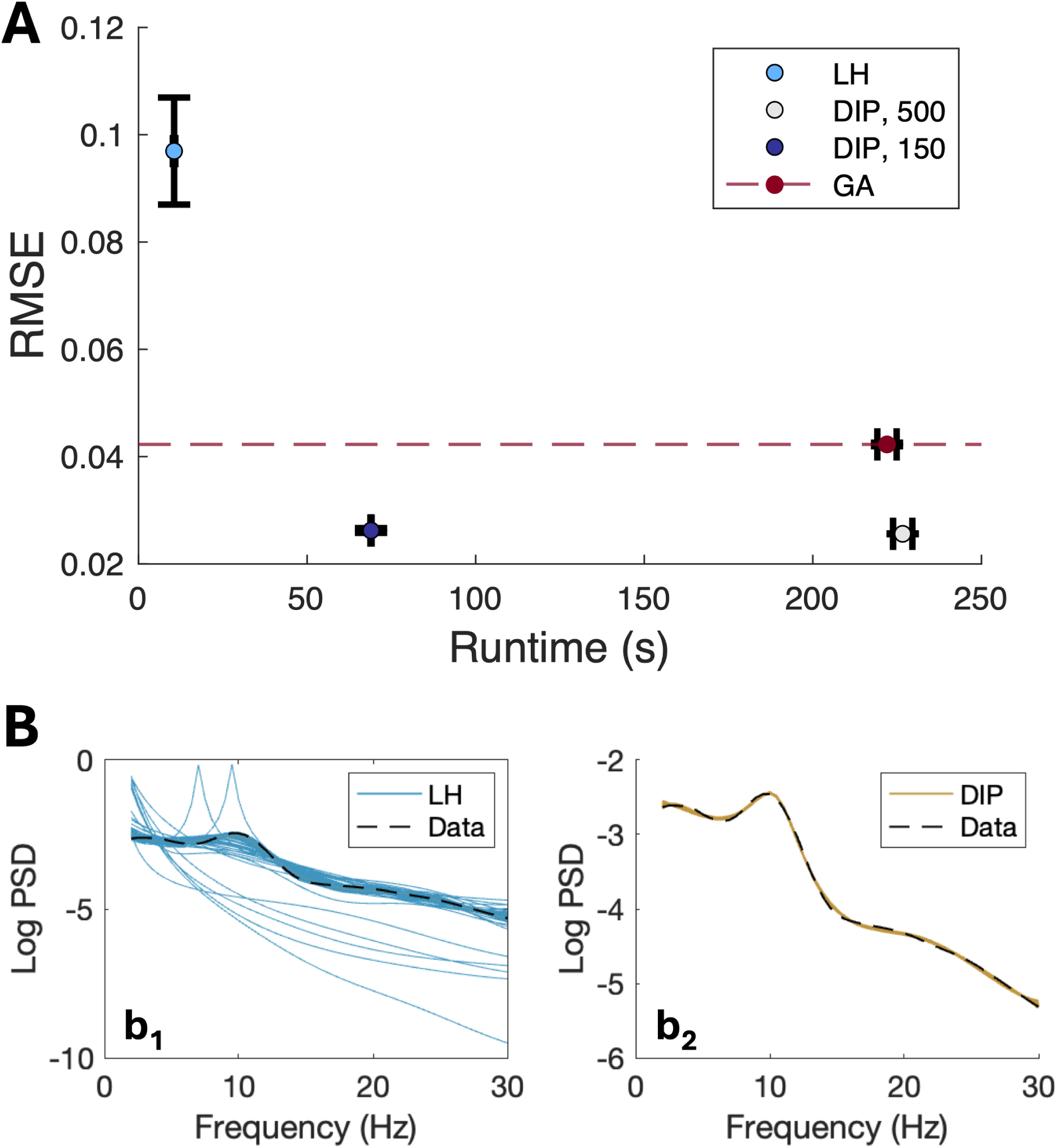
Efficiency of DIP-DCM. **A**: Reduction in the number of generations used in the first step of DIP-DCM. RMSE is the root mean square error between spectra of model and data, runtime is for a single inversion. Data are shown as mean S.E.M. The model was fitted to the PSD of the placebo condition, which serves as the critical baseline for all comparisons. **B**: Exemplar pre-placebo spectra of the model plotted against data. Model PSDs were obtained from randomly sampled priors (LH, *b*_1_) or dynamics-informed priors (DIP, *b*_2_). *Abbreviations*: LH, Latin hypercube; GA, genetic algorithm; gen, generations.

Moreover, it was also important to understand whether improvements similar to those of DIP-DCM could be achieved by seeding prior means at random via Latin Hypercube (LH), rather than generating DIP via GA. Prior covariance and bounds for LH-based models were identical to those of the DIP-DCM approach. While the LH-based approach was substantially faster, it failed to generate appropriate model dynamics (**Figure 5A,B**). Specifically, LH-based models exceeded the fitness threshold established by GA (**Figure 5A**), resulting in a majority of implausible models (**Figure 5B** *b*_1_), compared to DIP-based models (**Figure 5B,** *b*_2_).

Taken together, these results demonstrate that DIP-DCM generates optimal calibrations efficiently, requiring minutes to achieve accurate parameter estimation. Moreover, these results show that the GA can be terminated earlier once the prior is appropriately positioned within the parameter space, allowing the DCM variational inference to be leveraged for rapid convergence.

### 3.2 Study 2 - Application of DIP-DCM to MEG data

Study 1 (**Results 3.1**) indicated that different parameter estimation approaches can give rise to different mechanistic inferences for the same NMM and data. While dynamic causal models rely on a single normally-distributed prior, DIP-DCM takes into account additional parameter regimes and produces more informative predictions. To corroborate these findings, analyses were repeated on the Shaw et al., 2020 dataset (**Figure 6A**, **Methods 2.1**), providing a practical demonstration of the wider applicability of DIP-DCM.

**Figure 6:**
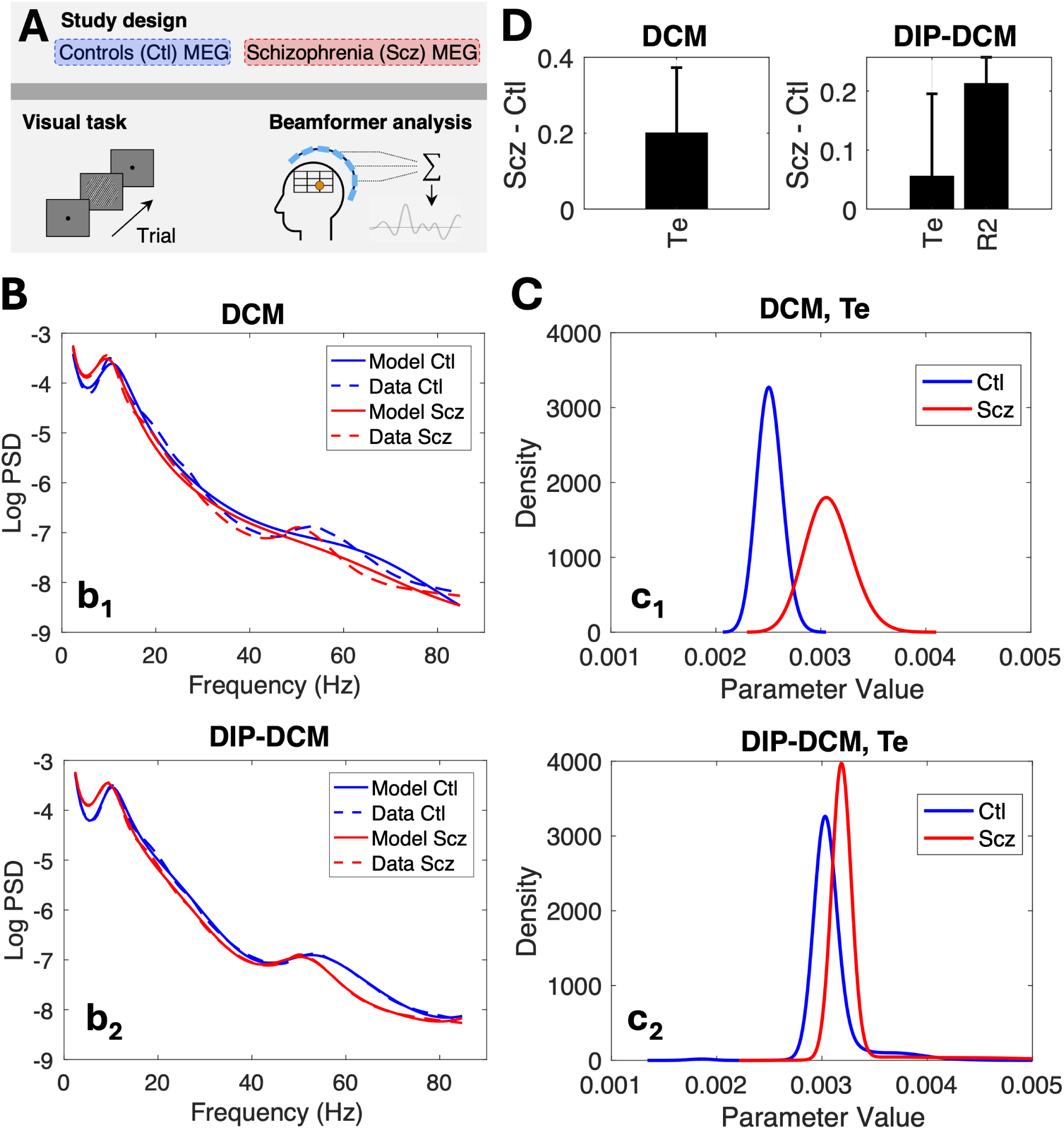
Application of DIP-DCM to MEG data. **A**: Study design. **B**: Data and model spectra generated by DCM (*b*_1_) and DIP-DCM(*b*_2_). Data is shown as mean and 95% CI.**C**: Representative examples of parameter distributions obtained via DCM (*c*_1_) and DIP-DCM (*c*_2_). All parameter densities are normalised to 1 and displayed within the same range. **D**: Mechanistic effects obtained as per **Methods 2.4**. *Abbreviations*: DIP-DCM, dynamic causal modelling with dynamics-informed priors; Ctl, control; Scz, schizophrenia.

Power spectra were modelled via DCM and the hybrid DIP-DCM workflow optimised for efficiency (see **Figure 5**; **Supplementary 3**). DIP-DCM yielded model dynamics that closely resembled data across the entire broadband spectrum (2 − 85 Hz), compared to DCM with “standard” priors (**Figure 6B**, panels *b*_1_, *b*_2_). This result is consistent with conclusions from **Results 4.2**.

Mechanistic inferences were obtained as per **Methods 2.4**, and examples of posterior distributions are shown in **Figure 6C** (DCM with “standard” priors in *c*_1_ and DIP-DCM in *c*_2_). Notably, the posterior distributions of parameter Te (**Figure 6C**) were largely uni-modal compared to those of study 1. For this parameter, both DIP-DCM and the standard DCM identified similar effects, indicating that impairments in visual processing and gamma oscillations in patients with schizophrenia are attributed to the speed of decay of excitatory activities (**Figure 6D**). However, DIP-DCM also identified an additional effect, namely an increase in parameter R2, which is indicative of higher firing thresholds and, therefore, of reduced responsiveness to sensory inputs. Similarly, in study 1 DIP-DCM detected key effects – such as the decrease in parameter Te following LEV administration – that DCM failed to identify. Taken together, these results indicate that DIP-DCM can help make disease-specific inferences with increased sensitivity.

## 4 Discussion

This paper compares the performance of three parameter estimation approaches to illustrate the challenge of parameter estimation in NMMs, where large and often degenerate parameter spaces hamper the inference process. While parameter priors in Bayesian inference are often informed by expert knowledge, this approach can bias the parameter estimation by overlooking alternative parameter configurations. Moreover, in cases where neuroscientific hypotheses remain unclear or the ‘true’ model is not represented among the candidate options, standard approaches may be limited. To address this problem, the present study introduces DIP-DCM, a dynamics-informed approach for DCM, designed to navigate complex parameter landscapes more effectively and to help draw inferences that are pure reflections of the information encoded in the data.

### 4.1 Reliability and validity of parameter inferences in DIP-DCM

Results from this study demonstrate that the choice of priors in DCM significantly impacts the validity of the posterior parameter estimates and the resulting inferences.

Typically, the challenge of finding suitable priors is addressed at the subject-level by using “empirical” or “data-informed” priors, where the *a priori* distributions for individual subjects are constrained by group-level estimates (Friston et al., 2015; Lee & Vanpaemel, 2018). A less common alternative involves deriving priors from a subset of individuals, effectively serving as a form of model re-training to narrow the parameter space for the remaining subset. However, these approaches may introduce bias by “overfitting” the model to the characteristics of the group, potentially overlooking inter-individual variability or the applicability of inferences to other cohorts. In addition, there may be cases where subject-level modelling may prove challenging, such as in studies with limited data, or where the goal is to fit directly the group average as, for example, in Tewarie et al., 2024 and in the present study. Thus, DIP-DCM is proposed as an alternative to guide the parameter inference process, starting from parameter values associated with optimal dynamics. Critically, rather than generating inferences that are fundamentally different from those of a standard DCM, DIP-DCM can identify additional effects with increased sensitivity, generating precise and testable hypotheses. This approach helps prevent biases introduced by a specific set of priors – which may lead to predictions that are not universally applicable – by considering a range of plausible outcomes. Thus, as the model parameters are often not uniquely identifiable from the data, their distributions may exhibit multiple peaks, or modes, each corresponding to a different parameter regime (Gäbor & Banga, 2015).

A detailed validation of the posterior estimates is provided in **Table 1** for key parameters. Within this study, parameters Te and Ti — representing the time scales of excitatory and inhibitory synaptic dynamics — are of particular importance. Different receptors are known to have different kinetics, determining the duration and nature of post-synaptic responses. While there is no direct mapping between parameter values and receptor types in this NMM, plausible receptor dynamics can be hypothesised based on the duration of the post-synaptic potential (PSP). Ultimately, the multi-modality of Te and Ti may reflect the dynamic nature of synaptic transmission and network activity. Each mode could correspond to distinct receptor kinetics, synaptic terminal properties, or features of excitatory post-synaptic currents (EPSCs) and inhibitory post-synaptic currents (IPSCs), as outlined in **Table 1**.

**Table 1:**
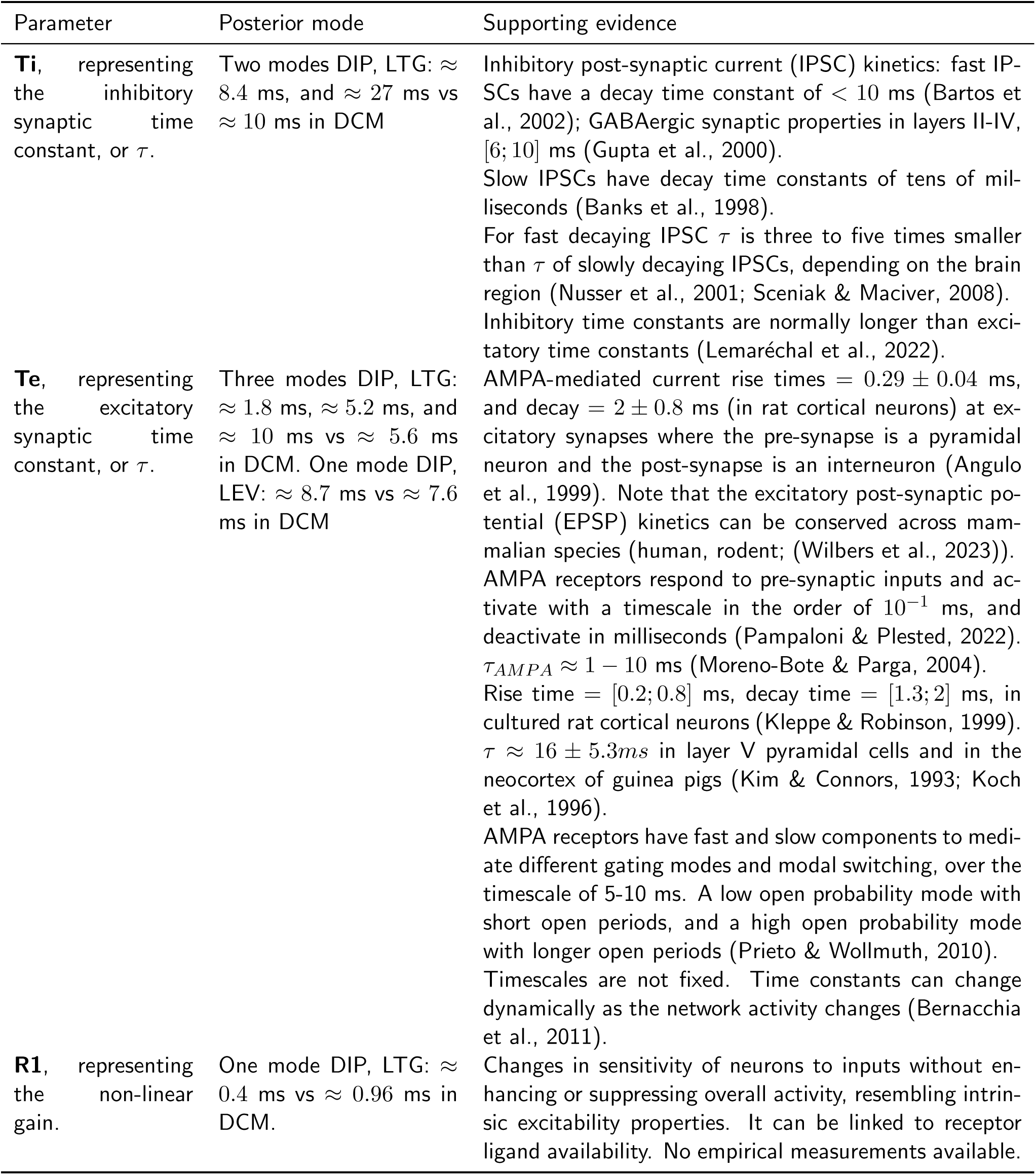
Posterior parameter modes generated by DIP-DCM in study 1.

The estimated parameters led to inferences presented in detail in **Discussion 4.2**. Acting as an intermediary between GA and DCM, DIP-DCM recovered key ASM-induced changes in model parameters, suggesting that both ASMs cause faster decays in excitatory activities, while LTG additionally slows the decay of inhibitory activities. Moreover, results suggest that LTG increases the parameter R1, indicating that small fluctuations in membrane potential produce larger changes in output firing rates. These inferences are in line with existing experimental evidence suggesting that LTG exerts a bidirectional modulation of both excitatory and inhibitory activity, while LEV appears to selectively affect excitatory activity (Greenhill & Jones, 2010; Margineanu & Klitgaard, 2003; Meehan et al., 2011).

Ultimately, the results support several important conclusions: (1) GA and DCM could produce qualitatively similar model dynamics, yet lead to different mechanistic inferences, which highlights the inherent challenges of parameter estimation in neural mass models. (2) DCM with “standard” priors is insufficiently sensitive to detect non-trivially small effects, such as the decrease in Te following LTG and LEV administration. (3) Global optimisations such as DIP-DCM prevent such type II errors, considering that the absence of an effect does not equate to the evidence of its absence. (4) DIP-DCM demonstrates both sensitivity and specificity, as evidenced by the effects of LEV on parameter Te and by the effects of schizophrenia on parameter R2.

### 4.2 Plausibility of DIP-DCM hypotheses on the effects of anti-seizure medication

DIP-DCM reconciled inferences on ASM effects across the tested approaches and identified the “true” group-level mechanistic fingerprints of ASMs for this NMM.

It is well established that LEV modulates presynaptic mechanisms involved in glutamate release at excitatory synapses, leading to changes in AMPA receptor-mediated currents (Carunchio et al., 2007). LEV binds to the presynaptic vesicle protein SV2A (Lynch et al., 2004; Sugaya et al., 2010), which has a modulatory role in transmitter release (Meehan et al., 2011). Consequently, LEV is unlikely to cause a substantial blockage of glutamatergic transmission (Meehan et al., 2011). The modulatory role of LEV aligns with inferences from DIP-DCM, indicating moderate effects on excitatory synapses. Moreover, the pharmacodynamics seem to involve a reduction in the release of vesicular content, which could explain the faster decay kinetics of EPSCs hypothesised from DIP-DCM. This is consistent with a theoretical framework where a reduction in neurotransmitter release may result in more rapid clearance, leading to faster decay of excitatory currents, or vice versa (Eggers & Lukasiewicz, 2006). This conclusion is also in line with studies conducted in epileptic rats showing that LEV increases the slope of field EPSPs (fEPSPs) (Salaka et al., 2022). Taken together, these results indicate that LEV only acts on excitatory neurons, supporting the hypothesis introduced by Margineanu and Klitgaard, 2003. These data also suggest that LEV may not simply suppress excitation but fine-tune and stabilise synaptic transmission towards healthier levels (Salaka et al., 2022).

However, different ASMs are known to act via different mechanisms (Rogawski & Löscher, 2004). ASMs are generally thought to modify the dynamic balance between excitation and inhibition, by either acting on excitation, inhibition, or on both concurrently. Differently from LEV, LTG has been suggested to increase the ratio between the global background synaptic excitation and inhibition (Greenhill & Jones, 2010). As indicated by DIP-DCM, this could plausibly occur through increasing the timescale separation between the excitatory and inhibitory synaptic currents. Nonetheless, earlier studies indicated a lack of observable changes in spontaneous EPSC and IPSC decays measured via patch-clamp following LTG administration (Cunningham & Jones, 2000). This highlights the complexity of relating cell-level recordings to properties of NMMs. Patch-clamp experiments, as those performed by Cunningham and Jones, 2000, may not accurately extrapolate population-level activity. In addition, this patch clamp experiment considered excitatory and inhibitory events in isolation, which may be inappropriate when investigating the effects of pharmacological agents that act across excitatory and inhibitory systems. Thus, if these results are ever to be tested empirically, it may be more appropriate to measure global background synaptic excitation and inhibition and synaptic plasticity as in Greenhill and Jones, 2010 and Salaka et al., 2022. Notably, the pronounced increase in inhibition-to-excitation ratio found by Greenhill and Jones, 2010 was paralleled by changes in neuronal excitability, which is supported by the changes in population sensitivity (parameter R1) inferred by DIP-DCM.

Taken together, these results show that DIP-DCM acts as a unifying framework, reconciling our understanding of the effects of pharmacological interventions across different experimental systems and providing hypotheses that can be readily tested.

### 4.3 Accelerated global search across dynamic causal models

This study highlights the advantages of using a hybrid DIP-DCM strategy, which enhances search efficiency while guaranteeing solution quality. Other two-phase hybrid optimisations — where global methods are used for broad sampling and local methods for rapid convergence — were shown to be very effective at addressing degeneracy and improving inference robustness across a wide range of optimisation problems. Hybrid optimisations, combining GAs and gradient descent-based (GD) methods, have proven useful for challenging problems with local minima outside the neuroimaging and DCM context (Gan et al., 2012), demonstrating superior performance in parameter estimation for biological network models (Rodriguez-Fernandez et al., 2006). Additionally, similarly to DIP-DCM, they have been shown to outperform GA alone in terms of efficiency and solution quality by positioning solutions in regions that facilitate convergence (D’Angelo & Palmieri, 2021). Performance remained superior even with a reduced number of generations, mirroring findings from the present study, where DIP-DCM achieved superior results for a third of the generations required by GA.

Furthermore, findings seem to resonate with similar strategies used within the DCM context, such as the massively parallel DCM (mpDCM) for hemodynamic models (Aponte et al., 2016). By adopting the concept of “thermodynamic integration” mpDCM computes multiple MCMC chains simultaneously at varying “temperatures”. Temperatures affect the trade-off between exploration and exploitation by adjusting the relative influence of the prior and likelihood in the sampling process, thereby guiding the trajectory from prior to posterior (Frässle et al., 2021; Aponte et al., 2022). This approach facilitated parameter exploration, preventing chains from being trapped in local optima, and accelerated convergence by allowing chains to exchange information through mechanisms inspired by evolutionary principles. While DIP-DCM differs from mpDCM in its mechanisms for parameter exploration and information exchange between search pathways, both methods share common goals. They improve convergence to solutions and adopt principled approaches to manage computational costs.

Finally, although alternative methods to variational Bayes have been used in the DCM framework for forward modelling of electrophysiological data — such as MCMC (Sengupta et al., 2015) — to the best of our knowledge this study provides the first publicly available tool where the DCM for M/EEG is driven by global optimisations.

### 4.4 Conclusions

To conclude, this study illustrates the existence of multiple calibrations, and associated inferences, for the same NMM and M/EEG data, and proposes a dynamics-informed framework for DCM as an alternative to traditional, local, inversion routines. This framework accurately estimates model parameters and the associated uncertainty by leveraging global optimisations, maintaining moderate computational costs, and suggesting potential benefits for a wider range of applications.

## Data and Code Availability

All analyses were conducted on data published previously by Biondi et al., 2022 and Shaw et al., 2020. Code used to generate figures including the database of parameter estimates can be found at: https://github.com/AlessiaCaccamo/Figures_DIP_DCM_25.git. The DIP-DCM toolbox code is open source, under the terms of the GNU General Public License, and can be found at: https://github.com/AlessiaCaccamo/DIP_DCM_25.git. This works on Windows, Linux, and macOS with an installed version of SPM12, and was not tested with other SPM versions. Code includes third-party functions (SPM, https://www.fil.ion.ucl.ac.uk/spm), with their respective copyright.

## Author Contributions

**AC**: Conceptualization, Methodology, Software, Validation, Formal analysis, Investigation, Data Curation, Writing - Original Draft, Writing - Review & Editing, Visualization, Project administration. **DMD**: Conceptualization, Methodology, Software, Resources, Writing - Review. **MPR**: Resources, Writing - Review. **ADS**: Conceptualization, Methodology, Resources, Writing - Review, Supervision. **MG**: Conceptualization, Methodology, Software, Resources, Writing - Review & Editing, Supervision. For the purpose of open access, the author has applied a Creative Commons Attribution (CC BY) licence to any Author Accepted Manuscript version arising from this submission

## Licence

For the purpose of open access, the authors have applied a Creative Commons Attribution (CC BY) licence to any Author Accepted Manuscript version arising from this submission.

## Acknowledgements

The authors wish to express their sincere gratitude to Prof. Karl Friston and Dr. Johan Medrano for their expert advice and insightful comments offered during a public presentation of this work. The authors also wish to thank Dr. Margaritis Voliotis and Dr. Kyle Wedgwood for their constructive feedback on early versions of the manuscript.

## Supplementary Material

The Supplementary material can be found here.

## Appendices

### A DCM with Dynamics Informed Priors (DIP-DCM)

Considering the set of optimal solutions *S_n_* generated by the GA (**Supplementary 2**), the set *S_m_* := {***θ*_1_**, ***θ*_2_***, …,* ***θ_m_***}, with *S_m_* ⊂ *S_n_*, was the set of *m* parameter vectors selected as follows:

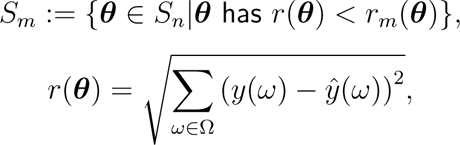

where *ŷ* and *y* are the model spectrum and the data spectrum at the frequency Ω, ranging between 2-30 Hz (for Biondi et al., 2022), or 2-85 Hz (for Shaw et al., 2020), *r_m_* is the *m^th^* smallest value of *r*(***θ***) over *S_n_*, and ***θ*** is the parameter vector ranked based on *r*(***θ***).

Thus, *S_m_* becomes the set of prior expectations for subsequent *m* VL inversions. Specifically, for each *m*-th parameter vector,

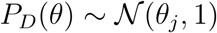

is the normally distributed dynamics-informed prior entering the DCM inversion routine.

This yields 500 normally distributed posterior distributions which are formalised into a probability density function as follows:

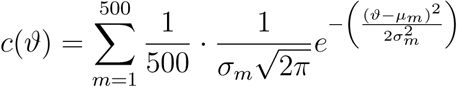

where *µ* and *σ*^2^ are the mean and variance of the posterior distribution.

### B Derivation of mechanistic inferences

Mechanistic inferences were extrapolated from the posterior parameter distributions for each of the three parameter estimation approaches.

For each *j^th^* parameter, 10^5^ random draws were taken from the posterior distributions of pre-treatment and post-treatment conditions, and denoted *z*(Post) and *z*(Pre), respectively. These distributions were the cumulative normalised sum of *m* normal distributions for DIP-DCM, the normal distribution for DCM, and the density function fitted to the distribution of GA estimates using the ksdensity MATLAB function.

Mean absolute differences were calculated as follows:

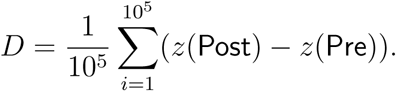

The respective 95% Bayesian credible intervals (BCIs) were calculated as follows, to determine the plausibility of nonzero differences:

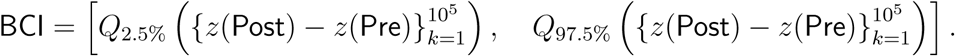

Intervals that excluded zero were interpreted as analogous to statistical significance at the *α* = 0.05 level.

Cohen’s d was also calculated to determine whether the study had sufficient power to support its conclusions,

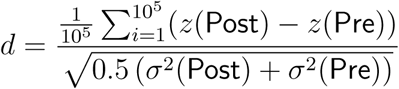

where *σ*^2^(Post), *σ*^2^(Pre) are the variances of *z*(Post) and *z*(Pre) samples.

Ultimately, the set *E* of effects *D* satisfying two criteria based on BCIs and Cohen’s d was the following:

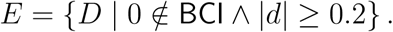

### C Parameter mapping and free energy landscapes

Free energy profiles were generated for each parameter of the Moran et al., 2007 model (**Methods 3.2**, **Appendix A**), to illustrate the challenges associated with parameter mapping in neural mass models. The approach was adapted from (Arand et al., 2015), considering the parameter contributions to the observed spectral response. A parameter was varied across a grid of 1001 values within biologically plausible bounds (**Supplementary 1**), which were centered around the standard priors. All other parameters were set to their “standard” SPM12 prior. Free energy was computed for each parameter combination via variational inversion, enabling exploration of the parameter space in the direction of maximum increase in free energy. Power spectra were calculated for each parameter regime to understand how the model dynamics change as a function of the priors and how this maps to free energy maximisation.

