## Supplementary Material for "Dynamics-Informed Priors (DIP) for Neural Mass Modelling"

### 1 Supplementary 1

#### 1.1 Neural mass equations

The model was from the SPM12 toolbox and is described in detail in Moran et al., 2007. It comprises three neuronal populations considered as single entities. The spiny stellate cell population, located in the granular layer, receives exeternal inputs, $I$, and excitatory inputs from pyramidal cells that are scaled by the connectivity parameter $G_1$. Their dynamics are formalised as follows:

$$\frac{dx_1}{dt} = x_4$$

$$\frac{dx_4}{dt} = \frac{H_e\left(\left(A_1 + A_3\right)S(x_9) + G_1 S(x_9) + I\right) - 2x_4 - \frac{x_1}{T_e}}{T_e}$$

where $x_1$ is the average post-synaptic potential (PSP) and $x_4$ the output signal of the population. $S(x)$ is the sigmoid transformation of the membrane potential into a firing rate, described as follows:

$$S(x) = \frac{1}{(1 + e^{-R_1(x-R_2)}) - (1 + e^{R_1 R_2})}$$

where $R_1$ and $R_2$ capture the slope and offset of the sigmoid function, respectively.

The excitatory pyramidal cell population, located in the infragranular layer, receives excitatory afferents from spiny stellate cells scaled by the connectivity parameter $G_2$, and inhibitory inputs scaled by the connectivity parameter $G_4$:

$$\frac{dx_2}{dt} = x_5$$

$$\frac{dx_5}{dt} = \frac{H_e\left(\left(A_2 + A_3\right)S(x_9) + G_2 S(x_1)\right) - 2x_5 - \frac{x_2}{T_e}}{T_e}$$

$$\frac{dx_3}{dt} = x_6$$

$$\frac{dx_6}{dt} = \frac{H_i G_4 S(x_{12}) - 2x_6 - \frac{x_3}{T_i}}{T_i}$$

$$\frac{dx_9}{dt} = x_5 - x_6$$

where $x_2$, $x_3$ are the PSPs, $x_5$, $x_6$ the firing rates, and $x_9$ the depolarisation depending on excitatory and inhibitory PSPs (EPSPs and IPSPs).

The inhibitory subpopulation, located in the supragranular layer, receives afferent input from the excitatory pyramidal cell population scaled by the connectivity parameter $G_3$. The inhibitory subpopulation also receives recurrent inhibitory inputs scaled by the connectivity parameter $G_5$. The associated state equations are as follows:

$$\frac{dx_7}{dt} = x_8$$

$$\frac{dx_8}{dt} = \frac{H_e\left(\left(A_1 + A_3\right) S(x_9) + G_3 S(x_9)\right) - 2x_8 - \frac{x_7}{T_e}}{T_e}$$

$$\frac{dx_{10}}{dt} = x_{11}$$

$$\frac{dx_{11}}{dt} = \frac{H_i G_5 S(x_{12}) - 2x_{11} - \frac{x_{10}}{T_i}}{T_i}$$

$$\frac{dx_{12}}{dt} = x_8 - x_{11}$$

where $x_7$, $x_{10}$ are the PSPs, $x_8$, $x_{11}$ the firing rates, $x_{12}$ the depolarisation depending on EPSPs and IPSPs.

Finally, $x_{13}$ plays a role attributable to a slow potassium conductance for the modulation of spike generation (Moran et al., 2007, 2013), as follows:

$$\frac{dx_{13}}{dt} = \frac{4S(x_1) - x_{13}}{T_k}.$$

The hidden neural state variables

$$\mathbf{x} = \begin{pmatrix} x_1 \\ x_2 \\ \vdots \\ x_{13} \end{pmatrix}$$

denoted by the vector $\mathbf{x} \in \mathbb{R}^{13}$, are mapped to the measured EEG data via the observation function:

$$g(\mathbf{x}) = f(\mathbf{x}, u, \boldsymbol{\theta}) + Bu.$$

where $f$ is the nonlinear neuronal model describing the evolution of the state variables over time, $\boldsymbol{\theta}$ is the set of parameters influencing how the states evolve, $g$ is the function that maps hidden neural states $x$ to the observations. $B$ is a vector describing the system's modulation by extrinsic inputs, and $u$ is the scalar input. Model parameters are reported in **Table S1**.

Table S1: **Parameter Priors and Bounds**. Prior parameter distributions were the default SPM12 priors, $\mu$ is the mean and $\sigma$ is the variance. The bounds on the parameters explored via the genetic algorithm (GA) are specified and reported here as [lower bound; upper bound]. Parameters are log-scaled to ensure positivity (Friston et al., 2019). Parameters $T_k$ and $H_i$ were fixed by default in the SPM model (see **Supplementary 1.4**).

| Parameters | DCM priors $P(\boldsymbol{\theta})$ | GA bounds |
|---|---|---|
| $R_1$ static nonlinearity, slope | $\mu = 1, \sigma = 0.1250$ | $[0.2431; 4.1132]$ |
| $R_2$ static nonlinearity, offset | $\mu = 2, \sigma = 0.1250$ | $[0.4862; 8.2264]$ |
| $T_e$ excitatory synaptic time constant | $\mu = 0.004, \sigma = 0.1250$ | $[0.000972; 0.0165]$ |
| $T_i$ inhibitory synaptic time constant | $\mu = 0.016, \sigma = 0.1250$ | $[0.0039; 0.0658]$ |
| $T_k$ potassium synaptic time constant | $\mu = 0.5120, \sigma = 0$ | $0.5120$ |
| $G_1$ pyramidal to stellate population connectivity | $\mu = 128, \sigma = 0.0625$ | $[46.8725; 347, 9401]$ |
| $G_2$ stellate to pyramidal population connectivity | $\mu = 128, \sigma = 0.0625$ | $[34.1660; 347, 9401]$ |
| $G_3$ pyramidal to inhibitory population connectivity | $\mu = 64, \sigma = 0.0625$ | $[23.4362; 183.0543]$ |
| $G_4$ inhibitory to pyramidal population connectivity | $\mu = 64, \sigma = 0.0625$ | $[23.4362; 174.0745]$ |
| $G_5$ inhibitory self-connectivity | $\mu = 4, \sigma = 0.0625$ | $[1.3521; 12.0371]$ |
| $H_e$ excitatory synaptic gain | $\mu = 8, \sigma = 0.0625$ | $[2.6449; 21.7463]$ |
| $H_i$ inhibitory synaptic gain | $\mu = 32, \sigma = 0$ | $32$ |
| $A_1$ extrinsic forward connectivity | $\mu = 32, \sigma = 0.5$ | $[1.2367; 541.3878]$ |
| $A_2$ extrinsic backward connectivity | $\mu = 16, \sigma = 0.5$ | $[0.9456; 312.2764]$ |
| $A_3$ extrinsic lateral connectivity | $\mu = 4, \sigma = 0.5$ | $[0.1510; 67.6735]$ |
| $D_e$ extrinsic propagation delay | $\mu = 0.002, \sigma = 0.0625$ | $[0.000736; 0.0054]$ |
| $D_i$ intrinsic propagation delay | $\mu = 0.016, \sigma = 0.0312$ | $[0.0079; 0.0348]$ |

#### 1.2 Spectral likelihood

Local linearity assumptions were adopted to circumvent the challenges of parameter estimation typical of nonlinear models and to enhance computational efficiency. The sigmoid function $S(x)$ is approximated via Taylor expansion around the steady state $\mathbf{x_0}$ using the defult SPM12 `spm_dcm_neural_x` function. At $x = 0$, the firing rate is described by:

$$S(\mathbf{x_0}) = \frac{1}{1+e^{R_1R_2}} - \frac{1}{1+e^{R_1R_2}} = 0,$$

meaning that all state variables have resting values of zero. The dynamics of the model around this point are dependent on the slope of the sigmoid function at $\mathbf{x_0}$,

$$\frac{dS}{dx} = \frac{R_1 e^{R_1R_2}}{(1+e^{R_1R_2})^2}.$$

The linear approximation of the system is:

$$\dot{x} = A\mathbf{x}(t) + Bu, \quad y_{obs} = C\mathbf{x}(t)$$

where

$$A_{i,j} = \frac{\partial \mathbf{f_i}}{\partial \mathbf{x_j}}\bigg|_{\mathbf{x}=\mathbf{x_0}}$$

is the Jacobian matrix representing the linear behaviour of the system around the steady state, and $C$ is a row vector mapping the hidden states to the observables, quantifying the contribution of the states $\mathbf{x}$ to the observed output $y_{obs}$. This mapping is described as follows:

$$C \in \mathbb{R}^{13}, \quad C_i = \begin{cases} 1, & \text{if } i = 9 \\ 0, & \text{if } i \neq 9 \end{cases}.$$

The Laplace transform was applied to the linear equations (Moran et al., 2007; Oppenheim et al., 1983) utilising the SPM12 `spm_csd_mtf` function to generate the power spectral density (PSD) of the model as follows:

$$Y(s) = C(sI - A)^{-1}B$$

where $s$ represents the complex Laplace variable, $A$ is the Jacobian matrix evaluated at $\mathbf{x_0}$, $C$ is the vector mapping the hidden states to the observed signal, $B$ is the input mapping, indicating how the inputs affect the system, and $I$ is the identity matrix.

This PSD is modulated by different sources of noise (endogenous neuronal fluctuations, measurement noise and data filtering that contributed to producing the observed PSD). This is conveyed by explicit parameterisation (Friston et al., 2015; Moran et al., 2009; Novelli et al., 2024; Razi et al., 2015). For details on the structure of this noise term, please refer to the section below (**Supplementary 1.3**). Ultimately, this spectral response is the predicted observation for the variational inversion and the GA optimisation.

The spectral likelihood, representing the probability of observing the PSD $y(\omega)$ for parameters $\boldsymbol{\theta}$ is defined as (Friston et al., 2012):

$$
\begin{aligned}
\log P(y \mid \boldsymbol{\theta}) = -\frac{1}{2}\sum_{\omega} \Big[ & \operatorname{Re}\big(y(\omega) - \hat{y}(\omega;\boldsymbol{\theta})\big)^{\top} \Pi_{\varepsilon} \operatorname{Re}\big(y(\omega) - \hat{y}(\omega;\boldsymbol{\theta})\big) \\
& + \operatorname{Im}\big(y(\omega) - \hat{y}(\omega;\boldsymbol{\theta})\big)^{\top} \Pi_{\varepsilon} \operatorname{Im}\big(y(\omega) - \hat{y}(\omega;\boldsymbol{\theta})\big) \Big] \\
& - \frac{1}{2}(\boldsymbol{\theta} - \boldsymbol{\theta_0})^{\top} \Pi_{\theta} (\boldsymbol{\theta} - \boldsymbol{\theta_0}) + \frac{1}{2}\log|\Pi_{\varepsilon}| + \frac{1}{2}\log|\Pi_{\theta}|
\end{aligned}
$$

where $y(\omega)$ is the PSD of the data at frequency $\omega$, $\hat{y}(\omega;\boldsymbol{\theta})$ is the PSD of the model at $\omega$, given the vector of parameters $\boldsymbol{\theta}$, $\boldsymbol{\theta_0}$ is the prior expectation, $\Pi_{\varepsilon}$ is the precision matrix of observation noise, $\Pi_{\theta}$ is the prior precision matrix over parameters (inverse of prior covariance).

#### 1.3 Observation noise

The observation function that generates the predicted spectral responses has two main contributions. One is neuronal, i.e., the pyramidal cell depolarisation, and the other is observation noise (**Supplementary 1.2**). Noise is modelled through parameterisation (**Table S2**; Friston et al., 2015; Razi et al., 2015; `spm_csd_mtf_gu` function from SPM12) of the observation function, modulating the predicted spectral response as follows:

$$Y_u(\omega) = e^{a_1} \cdot \omega^{-e^{a_2}} \cdot e^{Y_d d}$$

$$Y_n(\omega) = e^{b_1} \cdot \omega^{-e^{b_2}}$$

$$Y_s(\omega) = e^{c_1} \cdot \omega^{-e^{c_2}}$$

$$Y_2 = \hat{y_L} \cdot \rho(Y_u)$$

$$Y_{\text{norm}} = \frac{Y_2}{\int Y_2 \, df}$$

$$Y_{\log} = \log(Y_{\text{norm}})$$

$$\hat{y_m} = Y_{\log} + Y_s + Y_n$$

where $\hat{y_L}$ is the PSD of the model obtained via the Laplace transform. The parameters $a_1, a_2, b_1, b_2, c_1, c_2, d$ serve as modulations of the the predicted PSD. Specifically, $a_1$ is the amplitude of neuronal fluctuations, or innovations, i.e., spontaneous noisy activity, $a_2$ is the exponent of these innovations, $b_1, b_2, c_1, c_2$ capture amplitude and exponent of non-specific and specific channel noise (noise from nearby sources or from the recording channel). Parameters $d$ (i.e.,$d_1, d_2, d_3, d_4$) represent additional neuronal noise. Thus, $y_u$, $y_n$ and $y_s$ are spectral modulation vectors generated from noise parameters, considering that $\omega$ represents a vector of frequency bins dependent on the PSD of data and that $Y_d$ parameterises variations in individual spectral bins. $Y_2$ is an intermediate PSD adjusted to account for noise, and obtained by multiplying the PSD $\hat{y_L}$ by the diagonal of the noise parameterisation matrix $Y_u$, where $\rho$ is the diagonal function. The PSD is also modulated by $v_\omega$, representing effects of filtering, which is formalised as follows:

$$v_\omega = e^{f_1 + \frac{f_2 \omega}{n}}, \quad \text{for } \omega = 1, 2, \ldots, n$$

$$h = \hat{y_m} \odot v_\omega,$$

$f_1, f_2$ are the parameters for data filtering, $f$ is the frequency-dependent filter, $n$ is the total number of frequency bins, $\odot$ denotes element-wise multiplication, and $h$ is the model output PSD, representing the observed (or measured) output recapitulating empirical data. Therefore, the predicted observation, dependent on the parameters $\boldsymbol{\theta}$ and entering the inversion scheme, is:

$$\hat{y} = h(\boldsymbol{\theta}) + \epsilon,$$

with $\epsilon$ being the noise.

Table S2: **Priors and bounds on noise parameters**. Prior parameter distributions were the default SPM12 priors for the model, $\mu$ is the mean and $\sigma$ is the variance of the normal distributions. The bounds on the parameters explored via the genetic algorithm (GA) are reported as [lower bound; upper bound]. Parameters were log-scaled to ensure positivity.

| Parameters | DCM priors | GA bounds |
|---|---|---|
| $a_1$ neuronal fluctuations, amplitude | $\mu = 1, \sigma = 0.0078$ | $[0.7022; 1.4242]$ |
| $a_2$ neuronal fluctuations, exponent | $\mu = 1, \sigma = 0.0078$ | $[0.5571; 1.4242]$ |
| $b_1$ channel noise (non-specific), amplitude | $\mu = 1, \sigma = 0.0078$ | $[0.6856; 1.4242]$ |
| $b_2$ channel noise (non-specific), exponent | $\mu = 1, \sigma = 0.0078$ | $[0.7022; 1.4279]$ |
| $c_1$ channel noise (specific), amplitude | $\mu = 1, \sigma = 0.0078$ | $[0.6856; 1.4242]$ |
| $c_2$ channel noise (specific), exponent | $\mu = 1, \sigma = 0.0078$ | $[0.7022; 1.4279]$ |
| $d_1, d_2, d_3, d_4$ neuronal fluctuations | $\mu = 1, \sigma = 0.0078$ | $[0.6953; 1.4242]$ |
| | | $[0.7022; 1.4495]$ |
| | | $[0.7022; 1.5543]$ |
| | | $[0.6702; 1.4242]$ |
| $f_1, f_2$ data filtering | $\mu = 1, \sigma = 0.0156$ | $[0.6065; 1.6487]$ |
| | | $[0.5156; 1.6487]$ |

#### 1.4 Full and reduced models

Two neuronal parameters (Tk and Hi) were fixed by default in the SPM12 LFP model (**Table S1**). To examine this assumption, parameter estimation was conducted on both the reduced and full (**Table S3**) models, using dynamic causal modelling with dynamics-informed priors (DIP-DCM), (**Figure S1**).

Full and reduced models were compared via Bayesian model comparison. Considering that models at within each group had equal probability, $1/m$ with $m = 1, ..., 500$, the group-level log model evidence was calculated as follows:

$$log \sum_{m=1}^{500} e^{F_m},$$

where $F_m$ is variational free energy of each $m^{th}$ model in the group. Thus, the log Bayes factor ($\log BF$) for Bayesian model comparison was:

$$\log BF = \log \sum_{m=1}^{500} e^{F_{reduced,m}} - \log \sum_{m=1}^{500} e^{F_{full,m}}.$$

$\log BF > 3$ corresponds to statistical significance at $\alpha = 0.05$ (since $e^3 = 20$, a difference of 3 in log evidence corresponds to a ratio of $20^{-1} = 0.05$). Therefore, the full and reduced models had the same evidence, as $\log BF \approx 0$ for all experimental conditions (**Table S4**). However, since the full model required substantially longer computation times to generate the same dynamics (see **Figure 3B** in the main paper), this paper adopted the default reduced model.

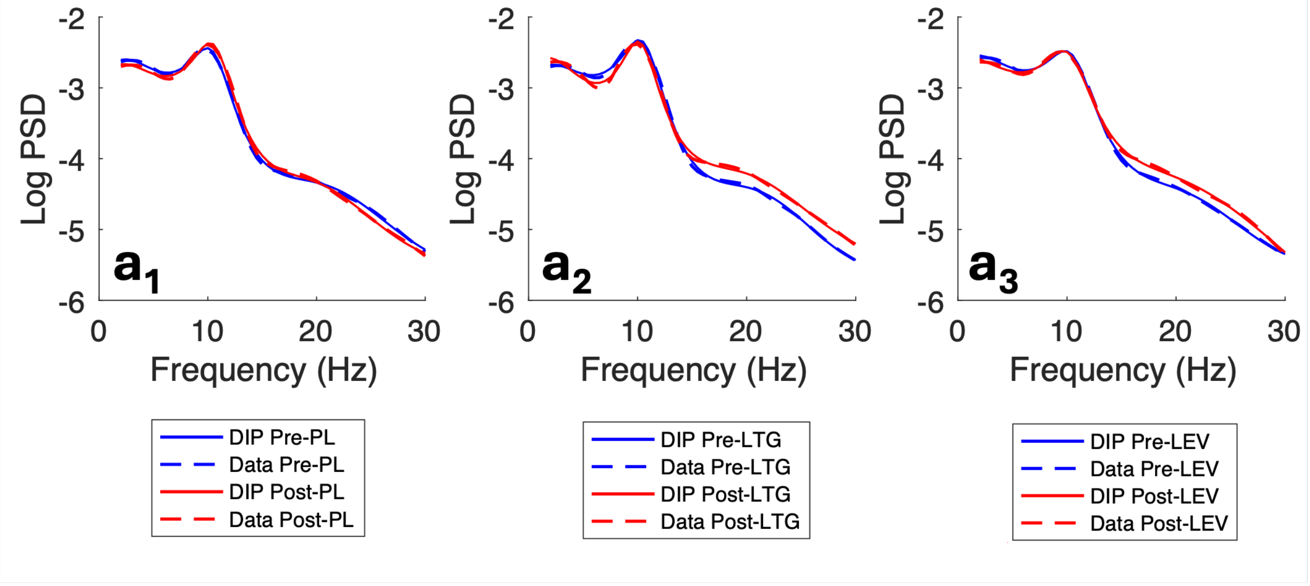


Figure S1: **Full LFP model**. Spectra of data and model for each experimental group: placebo (PL, $a_1$), lamotrigine (LTG, $a_2$) and levetiracetam (LEV, $a_3$). Data was from Biondi et al., 2022. *Abbreviations*: DIP, dynamics-informed priors.

Table S3: **Parameter priors and bounds for the full LFP model**. Parameter priors were the default SPM12 priors, $\mu$ is the mean and $\sigma$ is the variance. Parameter bounds utilised for the genetic algorithm (GA) are reported as [lower bound; upper bound] and are biologically realistic.

| Parameters | DCM priors | GA bounds |
|---|---|---|
| $R_1$ static nonlinearity, slope | $\mu = 1, \sigma = 0.1250$ | $[0.2431; 4.1132]$ |
| $R_2$ static nonlinearity, offset | $\mu = 2, \sigma = 0.1250$ | $[0.4862; 8.2264]$ |
| $T_e$ excitatory synaptic time constant | $\mu = 0.004, \sigma = 0.1250$ | $[0.000972; 0.0165]$ |
| $T_i$ inhibitory synaptic time constant | $\mu = 0.016, \sigma = 0.1250$ | $[0.0039; 0.0658]$ |
| $T_k$ potassium synaptic time constant | $\mu = 0.5120, \sigma = 0.125$ | $[0.1017, 2.1060]$ |
| $G_1$ pyramidal to stellate population connectivity | $\mu = 128, \sigma = 0.0625$ | $[46.8725; 347, 9401]$ |
| $G_2$ stellate to pyramidal population connectivity | $\mu = 128, \sigma = 0.0625$ | $[34.1660; 347, 9401]$ |
| $G_3$ pyramidal to inhibitory population connectivity | $\mu = 64, \sigma = 0.0625$ | $[23.4362; 183.0543]$ |
| $G_4$ inhibitory to pyramidal population connectivity | $\mu = 64, \sigma = 0.0625$ | $[23.4362; 174.0745]$ |
| $G_5$ inhibitory self-connectivity | $\mu = 4, \sigma = 0.0625$ | $[1.3521; 12.0371]$ |
| $H_e$ excitatory synaptic gain | $\mu = 8, \sigma = 0.0625$ | $[2.6449; 21.7463]$ |
| $H_i$ inhibitory synaptic gain | $\mu = 32, \sigma = 0.0625$ | $[11.7721; 91.7196]$ |
| $A_1$ extrinsic forward connectivity | $\mu = 32, \sigma = 0.5$ | $[1.2367; 541.3878]$ |
| $A_2$ extrinsic backward connectivity | $\mu = 16, \sigma = 0.5$ | $[0.9456; 312.2764]$ |
| $A_3$ extrinsic lateral connectivity | $\mu = 4, \sigma = 0.5$ | $[0.1510; 67.6735]$ |
| $D_e$ extrinsic propagation delay | $\mu = 0.002, \sigma = 0.0625$ | $[0.000736; 0.0054]$ |
| $D_i$ intrinsic propagation delay | $\mu = 0.016, \sigma = 0.0312$ | $[0.0079; 0.0348]$ |

Table S4: **Bayesian model comparison between the reduced and full model**. For each of the six sub-datasets (pre- and post-PL, LTG, LEV) from Biondi et al., 2022, the reduced model had variational free energy $F_{reduced}$, and the full model had variational free energy $F_{full}$. Each model was a group of $m$ models, as defined by the respective dynamics-informed priors. *Abbreviations:* PL, placebo; LTG, lamotrigine; LEV, levetiracetam.

| Dataset | $F_{reduced} - F_{full}$ |
|---|---|
| $Pre - PL$ | $0.1286$ |
| $Post - PL$ | $-0.0442$ |
| $Pre - LTG$ | $-0.1294$ |
| $Post - LTG$ | $-0.5011$ |
| $Pre - LEV$ | $0.1926$ |
| $Post - LEV$ | $0.3210$ |

### 2 Supplementary 2

#### 2.1 Dynamic causal modelling

Prior beliefs on parameter values (**Table S1**) entered the traditional VL routine to obtain an updated posterior parameter density and a free energy approximation to model evidence (see Friston et al., 2003; Parr et al. T. and Friston, 2022; Zeidman et al., 2023 for in-depth derivation). By Bayes' product rule, the true posterior is:

$$P(\boldsymbol{\theta}|y) = \frac{P(y|\boldsymbol{\theta})P(\boldsymbol{\theta})}{P(y)},$$

where $y$ is the PSD of the data used to infer the parameters $\boldsymbol{\theta}$, $P(\boldsymbol{\theta})$ are the prior beliefs, $P(y|\boldsymbol{\theta})$ is the likelihood, and $P(y)$ is model evidence, or marginal likelihood. Considering that the likelihood $P(y|\boldsymbol{\theta})$ cannot be calculated, the true posterior distribution $P(\boldsymbol{\theta}|y)$ is approximated with a simpler distribution $Q(\boldsymbol{\theta})$, which, under the Laplace assumption, is considered to be Gaussian and specified as expectation (mean) and variance. The marginal likelihood, or model evidence, is approximated by free energy, defined as follows:

$$F[Q, y] = D_{KL}[Q(\boldsymbol{\theta})||P(\boldsymbol{\theta})] - E_Q(\boldsymbol{\theta})[\ln P(y|\boldsymbol{\theta})],$$

$$F[Q, y] \approx P(\boldsymbol{\theta}),$$

where free energy $F[Q, y]$ is a functional of the approximate posterior, $Q$, and of the data $y$. The first term represents model complexity and it is quantified as Kullback–Leibler (KL) divergence ($D_{KL}$) between the approximate posterior $Q(\boldsymbol{\theta})$ and the prior $P(\boldsymbol{\theta})$. The second term is accuracy and it is the expected value with respect to the variational distribution $Q(\boldsymbol{\theta})$, ($E_Q(\boldsymbol{\theta})$), of the log likelihood ($\log P(y|\boldsymbol{\theta})$).

Thus, parameter estimation is treated as an optimisation problem, where parameters are optimised in the direction of maximum increase of $F$.

#### 2.2 Genetic algorithm

The genetic algorithm (GA) was adapted from Dunstan et al., 2023 and utilised the NSGA-II variant from the MATLAB `gamultiobj` function (Deb, 2001). The algorithm initialises a population of parameter vectors $V$, representing solutions to the optimisation problem, defined as:

$$V(t) = \{\boldsymbol{\theta_1}, \boldsymbol{\theta_2}, \ldots, \boldsymbol{\theta_p}\},$$

where $p$ is a population of parameter sets, and $p = 650$. The population $V$ evolves for $t$ generations, with $t = (0, 1, ..., 500)$ for study 1 and $t = (0, 1, ..., 150)$ for study 2. Each parameter set is a vector $\boldsymbol{\theta}$ of 27 parameter values $\theta_j$. The initial population is randomly sampled via Latin hypercube as described in (Dunstan et al., 2023), within specified bounds $[lb_j, ub_j]$ (**Table S1**).

The algorithm utilised two objective functions, and thus it can be referred to as a multi-objective GA. The objectives $J_1$ and $J_2$ were evaluated on each parameter set in the population $V(t)$ at each generation $t$, as follows:

$$J_1 = \sqrt{\sum_{\omega \in \Omega_1} (y(\omega) - \hat{y}(\omega))^2}$$

$$J_2 = \sqrt{\sum_{\omega \in \Omega_2} (y(\omega) - \hat{y}(\omega))^2}$$

where $\hat{y}$ is the PSD of the model, $y$ is the PSD of the data at the frequencies $\Omega_1$, $\Omega_2$ ranging between 6.5-12.5 Hz, and 11.5-30 Hz, respectively for the Biondi et al., 2022 dataset. These objectives ensure that the alpha and beta frequencies are well-represented. A similar logic was applied to the Shaw et al., 2020 dataset, with $\Omega_1$, ranging between $[10.7 - 65.8]$ Hz and $\Omega_2$ ranging between $[35.6 - 85]$ Hz.

Thus, Parameter sets are selected from the population based on their fitness scores, determined by the values of each objective. The built-in `selection tournament` selection function randomly chooses groups of four individuals from the current population. The individual with the best fitness score in the group is chosen as a "parent" to produce offspring for the next generation. Offspring is produced via crossover and mutation (`crossover scattered` and `mutation adapt feasible` functions from the matlab toolbox), evolving the population towards increasingly better solutions. Solution rank is measured by Pareto dominance:

$$\boldsymbol{\theta_A} \in P(t_{end}) \text{ Pareto dominates } \boldsymbol{\theta_B} \in P(t_{end}) \text{ if}$$
$$\forall k = 1, 2, f_k(\boldsymbol{\theta_A}) \leq f_k(\boldsymbol{\theta_B}) \text{ and } \exists k = 1, 2 : f_k(\boldsymbol{\theta_A}) < f_k(\boldsymbol{\theta_B}),$$

with $P(t_{end})$ the set of all solutions. This generated a Pareto front of solutions, i.e., a set of points in the 2D objective space where no solution can be improved without worsening the other. From the Pareto front one solution is selected — the solution with smallest Euclidean distance between the two objectives in the objective space. The GA is repeated $n$ times, with $n = 1000$, yielding a set of solutions:

$$S_n := \{\boldsymbol{\theta_1}, \boldsymbol{\theta_2}, \ldots, \boldsymbol{\theta_n}\}.$$

### 3 Supplementary 3

The performance the DIP-DCM parameter estimation approach was assessed via two complementary analyses (**Figure S2**). These analyses were performed using data from the placebo condition of study 1 (Biondi et al., 2022), serving as a baseline comparison.

First, the number of generations used in the global search – implemented in DIP-DCM as a genetic algorithm – was systematically reduced to determine the minimum required to produce stable model dynamics. Notably, 150 generations seemed to be necessary to initialise the DCM variational inference sufficiently close to the optimum and achieve optimal model fitness, as measured via root mean square error (RMSE). This configuration also maintained moderate computation times for each individual inversion (**Figure S2A**). Notably, DIP-DCM outperformed a genetic algorithm alone as well as a DCM initiated from randomly sampled priors using a Latin Hypercube (LH), highlighting the best trade-off between accurate estimation and computational efficiency.

Second, with the number of generations fixed at 150, the number of priors, and therefore the number of variational inversions, was systematically reduced from 500 to 50 (**Figure S2B**) and selected based on the RMSE fitness scores. Notably, when the number of priors was reduced below 400, there was a gradual decline in parameter inference accuracy, with spurious effects beginning to emerge at 100 and 50 inversions. Mechanistic effects were calculated from the estimated parameter distributions as per **Methods 3.4**. Results suggest that a sufficiently large number of inversions is required to average out noise and obtain statistically reliable estimates. However, it is also possible that improved methods for selecting informative priors could mitigate these effects.

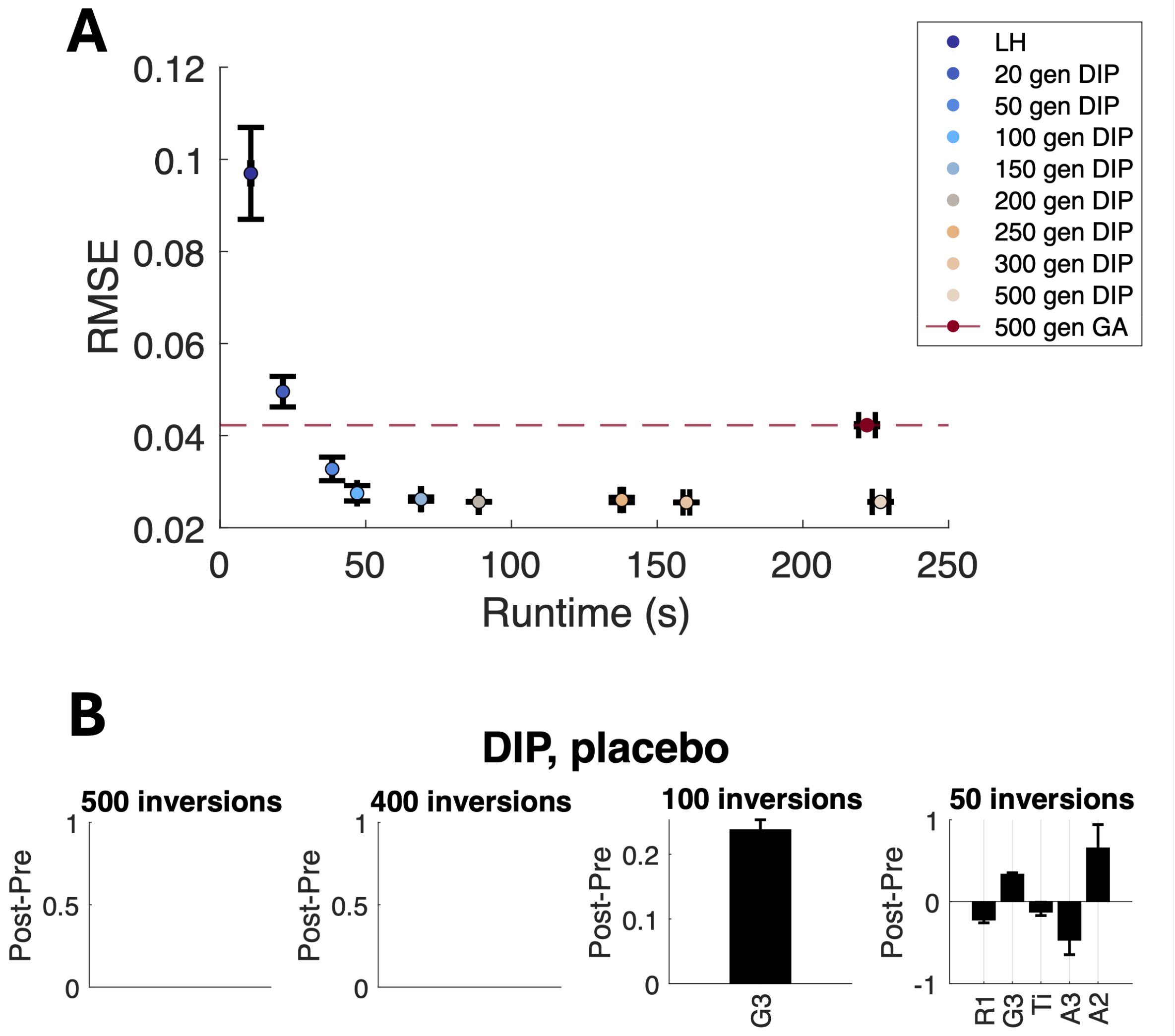


Figure S2: **Efficiency of the dynamics-informed approach**. **A**: Systematic reduction in the number of generations used by the genetic algorithm (GA). RMSE is the root mean square error between spectra of model and data for the pre-placebo experimental group, runtime is for each GA or DIP-DCM inversion (n=500). Data is shown as mean $\pm$ S.E.M. **B**: With the number of generations fixed at 150, the number of priors was systematically reduced from 500 to 50 in steps of 50. Placebo effects were calculated using the estimated parameter distributions as per **Methods 3.5** and shown as mean differences and $95\%$ bayesian credible intervals. *Abbreviations*: PL, placebo; gen, generations; DIP-DCM, dynamic causal modelling with dynamics-informed priors; LH, Latin hypercube.
